# Recurrent innovation of protein-protein interactions in the *Drosophila* piRNA pathway

**DOI:** 10.1101/2024.10.05.616775

**Authors:** Sebastian Riedelbauch, Sarah Masser, Sandra Fasching, Sung-Ya Lin, Harpreet Kaur Salgania, Mie Aarup, Anja Ebert, Mandy Jeske, Mia Levine, Ulrich Stelzl, Peter Andersen

## Abstract

Despite being essential for fertility, genome defence pathway genes often evolve rapidly. However, little is known about the molecular basis of this adaptation. Here, we characterize the evolution of a protein interaction network within the PIWI-interacting small RNA (piRNA) genome defence pathway in *Drosophila* at unprecedented scale and evolutionary resolution. We uncover pervasive rapid evolution of a protein interaction network anchored at the Heterochromatin Protein 1 (HP1) paralog Rhino. Using complementary phylogenetic analysis, high-throughput yeast-two-hybrid matrix screening, and *in vivo* interaction analyses in cross-species transgenic flies, we characterized three distinct evolutionary protein interaction trajectories across ∼40 million years of *Drosophila* evolution. The data set covering 11 piRNA pathway proteins of five *Drosophila* species revealed several protein interactions that are fully conserved, indicating functional conservation despite overall rapid amino acid sequence change. Other interactions are preserved through co-evolution and were detected only between proteins within or from closely related species. We also identified sets of species-restricted protein interactions which, through rewiring of a Rhino-anchored transcription factor network, may preserve critical roles in enabling and adapting piRNA production from heterochromatic loci. In sum, our analyses dissected principles of interaction evolution in an adaptively evolving protein-protein interaction network uncovering evolutionary and functional insight into germline piRNA production across *Drosophila* species. Our work provides key experimental evidence in support of a model proposing that intermolecular interaction innovation is a major molecular mechanism of evolutionary adaptation in protein-coding genes.

## INTRODUCTION

The patterns of adaptive evolution, also referred to as positive selection, across protein-coding genes make strong predictions about the molecular processes driving such innovation. Specifically, amino acids located at the surface of folded protein domains – and thus available to make intermolecular interactions – evolve faster than buried residues not exposed to the surrounding solvent^1^. Consistently, amino acids in protein termini and intrinsically disordered regions, which are often solvent exposed, evolve faster than central and ordered regions, respectively, and are often targeted by adaptive evolution^1–4^. Furthermore, large-scale comparison of protein architecture, functional annotation, and sequence evolution suggest that adaptive protein evolution is mainly driven by intermolecular interactions, such as protein-protein interactions^1,5^. A few studies have examined the evolution of protein-protein interactions between yeast species^6^ or across vast evolutionary time between yeast and humans^7^. However, we lack studies with the stepwise granularity in evolutionary time needed to connect to the molecular innovation step in the studied proteins. Specifically, investigations comparing only two species captures only one possible evolutionary trajectory. Other evolutionary outcomes are left undiscovered, limiting our grasp of the diverse mechanisms of sustaining an essential pathway under evolutionary pressure to change. Large-scale examination of protein-protein interactions in adaptively evolving protein interaction networks therefore remains outstanding.

The PIWI-interacting RNA (piRNA) pathway is a major transposable element (TE) silencing mechanism in gonads of most animals. Here, PIWI-clade Argonaute proteins guided by their bound piRNAs induce transcriptional and post-transcriptional silencing of target transposable elements^8–13^. While the piRNA pathway is present in most animals, the piRNA source loci, biogenesis mechanisms, and regulatory targets differ between model organisms and developmental stages. Furthermore, functional diversification of the piRNA pathway has taken place in the recent evolutionary history of multiple animal species, including several *Drosophila* species^14^ and teleost fish^15^. In *Drosophila*, key genes of the piRNA pathway evolve rapidly under positive selection, including genes involved in piRNA precursor synthesis^16^, processing^17,18^, and in piRNA-mediated silencing^17–20^. Furthermore, factors assumed to be essential for piRNA-mediated silencing based on studies using *D. melanogaster* have been found to be dispensable for transposon silencing in closely related species^21^. Given the experimental tractability of the *D. melanogaster* model organism, the *Drosophila* piRNA pathway represents a tractable model system to investigate the molecular mechanistic diversification resulting from rapid evolution within animal protein interaction networks.

Amongst the fastest evolving piRNA pathway genes in *D. melanogaster* are several involved in germline piRNA precursor biogenesis^16,22^. Most germline piRNAs in *Drosophila* are produced from heterochromatic loci and therefore require a dedicated transcription and export machinery for their expression. At the base of this mechanism is the HP1 homolog protein, Rhino (*rhi*)^23^, which is recruited to piRNA source loci through binding to heterochromatic Histone 3 trimethyl-modifications at Lysine 9 (H3K9me3) and Lysine 27 (H3K27me3)^24–26^ as well as the sequence-specific zinc finger protein, Kipferl (*Kipf*)^27^. Via the adaptor protein Deadlock (Del), Rhino in turn recruits a network of effector proteins that enable the production of piRNA precursors from the heterochromatic piRNA source loci (**Figure 1A**). First, an alternative basal transcription factor complex consisting of the Transcription-factor-IIA (TFIIA) homolog, Moonshiner (Moon), TFIIA-S and TATA box binding protein-related factor 2 (Trf2) is recruited to initiate heterochromatin-dependent piRNA precursor transcription^28^, while canonical activation of transposon promoters is thought to be suppressed by the C-terminal Binding Protein (CtBP) transcriptional repressor^29^. Following, co-transcriptional splicing as well as cleavage and polyadenylation is suppressed via recruitment of the DXO-like protein Cutoff (Cuff)^24,30–32^. Finally, piRNA precursor export is mediated by an alternative RNA export complex containing Nuclear export factor 3 (Nxf3), NTF2-related export protein 1 (Nxt1) and Bootlegger (Boot), which further connect to the general mRNA export complex THO via the UAP56 protein^33–36^. While these factors are all essential for transposon control and female fertility in *D. melanogaster*, several observations suggest that proteins enabling germline piRNA precursor production undergo rapid adaptation. For example, ovaries of the progeny of hybrid crosses between the closely related species *D. melanogaster* and *D. simulans* show elevated TE activity and reduced piRNA precursor production from dual-strand clusters, indicating that the sequence divergence in piRNA precursor biogenesis genes has resulted in functional divergence between the two species^37^. Furthermore, two studies found that the *D. simulans* orthologs of Rhino, Deadlock and Cutoff proteins are not able to rescue the respective ortholog mutants in *D. melanogaster*, likely due to divergence in the protein-protein interaction surfaces connecting Rhino, Deadlock and Cutoff^29,38,39^. These findings suggest that recurrent molecular innovation, likely at the level of protein-protein interactions, is required to maintain efficient germline transposon silencing in *Drosophila*. However, how adaptive evolution has diversified the proteins but preserved the functional output of piRNA biogenesis remains unknown.

**Figure 1.**
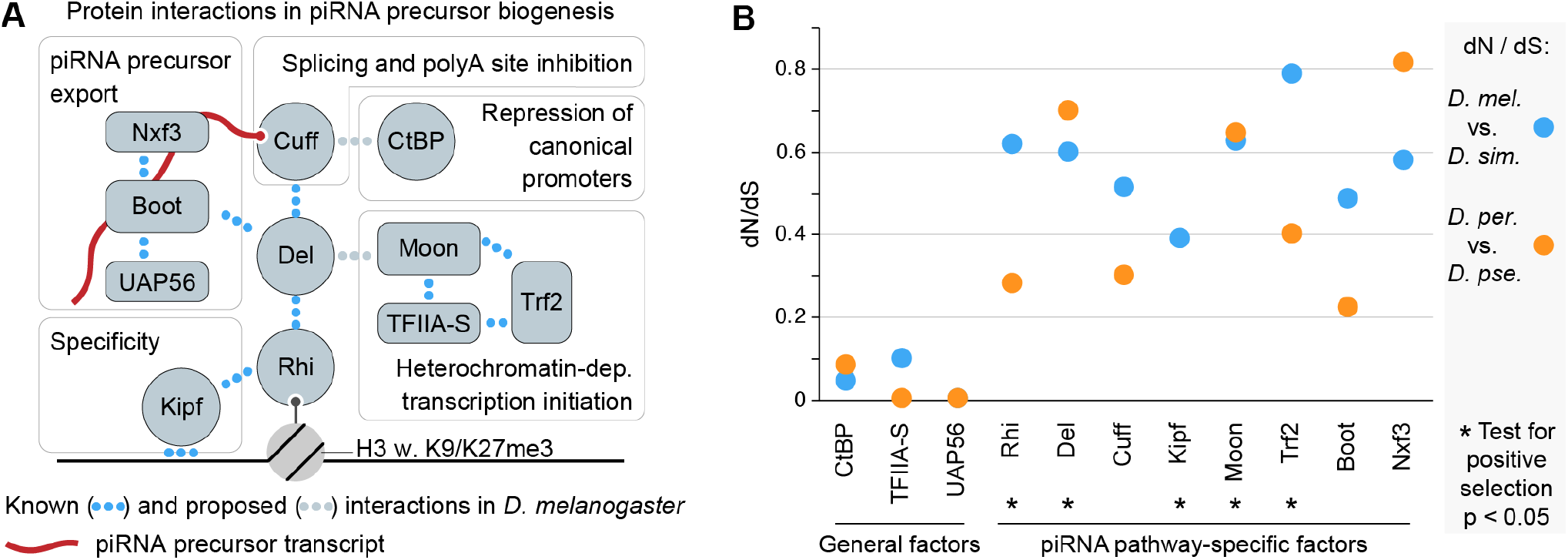
Rapid evolution of the piRNA biogenesis protein network. (A) Schematic showing protein interactions amongst factors enabling germline piRNA precursor biogenesis. Interactions validated to be direct are shown in blue dots, while proposed interactions are shown in grey. (B) Plot showing dN/dS ratios of genes encoding the proteins shown in (A). Ratios were calculated between *D. melanogaster* and *D. simulans*. Genes with significant (p < 0.05) codeml results from PAML suggesting positive selection are indicated by black stars.

Here, we comprehensively examine the evolution of protein-protein interactions within the piRNA precursor biogenesis pathway in five *Drosophila* species, representing spans of 2 to 40 million years of evolution. To do so, we performed high-throughput yeast-two-hybrid (Y2H) matrix interaction screening with 11 proteins within and between species, complemented by molecular evolution analysis, *Drosophila* cell culture assays, and *in vivo* cross-species co-immunoprecipitations (coIPs) from ovaries. We find that the overall interaction network structure of piRNA precursor biogenesis proteins, despite pervasive signatures of rapid, adaptive evolution, is conserved across the investigated species. However, our data also reveals a highly heterogenous mix of individual protein interactions between piRNA precursor biogenesis factors, including conserved interactions (e.g. Del– Cuff), protein-protein co-evolution (e.g. Del–Boot), as well as species-restricted innovation in the protein interaction network by “rewiring” that alters how transcription factors acting at germline piRNA source loci are recruited. Thus, by simply asking the central question of protein-protein interaction compatibility amongst different species, this study shows how rapid evolution has shaped a protein network under positive selection and reveals substantial mechanistic diversity in piRNA precursor production between even closely related species. We speculate that such biochemical innovation reshapes the expressed piRNA population to counteract threats to genome integrity.

## RESULTS

### Rapid evolution and positive selection in genes mediating piRNA precursor biogenesis

Evidence of positive selection or patterns of rapid evolution have been reported for several genes in the Drosophila piRNA pathway. However, a broader coherent molecular evolution analysis, including more recently discovered factors, is missing for a comprehensive understanding of the evolutionary history of the piRNA pathway. To fill this gap, we performed molecular evolution analyses of factors required for germline piRNA precursor biogenesis in *D. melanogaster* (**Figure 1A**). This piRNA precursor production relies on highly specialized piRNA pathway genes as well as deeply conserved genes that serve general functions in RNA polymerase II-based gene expression, including TFIIA-S, CtBP, and UAP56. Deeply conserved functions are expected to be reflected in very low frequency of amino acid-altering (non-synonymous) mutations, thereby providing a within-pathway control group. Indeed, comparing coding sequences between two closely related Drosophila species pairs (*D. melanogaster* to *D. simulans* and *D. persimilis* to *D. pseudoobscura*), dN/dS (ratio of nonsynonymous to synonymous divergence) estimates for all tested piRNA pathway-specific factors was notably elevated compared to the three genes also serving more general roles in gene expression (**Figure 1B**). Of note, for Trf2, which functions both in the piRNA pathway^28^ and in canonical gene expression^40,41^, we observed high sequence divergence concentrated in the long isoform, for which the function remains unresolved. Rapid sequence divergence can be explained by either genetic drift (lack of functional constraint) or by adaptive evolution (positive selection for new sequence variants). To determine the extent of adaptive evolution within the germline piRNA precursor biogenesis network, we performed likelihood ratio tests between species of the *melanogaster* species group, spanning ∼25 million years of divergence^42^. We inferred a history of adaptive evolution for Rhino, Deadlock, Kipferl, Moonshiner, and Trf2 (**Figure 1B, Figure S1A**, and **Table S1**). In sum, we extend previous observations of rapid evolution within the piRNA pathway to uncover pervasive rapid, adaptive evolution in genes involved in germline piRNA precursor production. These observations suggest recurrent functional innovation within the essential protein interaction network supporting *Drosophila* piRNA biogenesis.

### A systematic Y2H screen defines the protein interaction network evolution of piRNA precursor biogenesis

Germline piRNA precursor biogenesis is facilitated by protein-protein interactions that lead to the recruitment of effectors (**Figure 1A**) driving transcription, co-transcriptional processing, and piRNA precursor export^24,28,31,35,36^. Interestingly, recent studies have uncovered incompatibilities amongst the factors Rhino, Deadlock and Cutoff from the closely related *D. melanogaster* and *D. simulans* species^29,38^. Given the recent predictions that adaptive evolution in protein-coding genes is mainly driven by intermolecular interactions rather than innovation in e.g. protein stability or folding^1,5^, we therefore hypothesized that the functional innovation suggested by our evolutionary analyses has occurred at the level of protein-protein interactions. To test this hypothesis, we performed a cross-species, all-versus-all yeast-two-hybrid (Y2H) matrix screen, a method previously used to study host-pathogen relationships (reviewed in refs. 43,44), protein interactome essentiality^45^, and subcomplex contacts in the nuclear pore complex^46^. Here we screened 11 proteins involved in germline piRNA precursor biogenesis **(Figure 1A)** from five different *Drosophila* species (*D. melanogaster, D. simulans, D. erecta, D. persimilis*, and *D. virilis*), representing different evolutionary distances across 40 million years of divergence (**Figure 2A**). As Kipferl is absent in *D. persimilis* and *D. virilis*^27^, this resulted in a set of 54 proteins from five species (**Table S2**) allowing a systematic assessment of 308 intra- and 1123 inter-species unique protein pairs for interaction.

**Figure 2.**
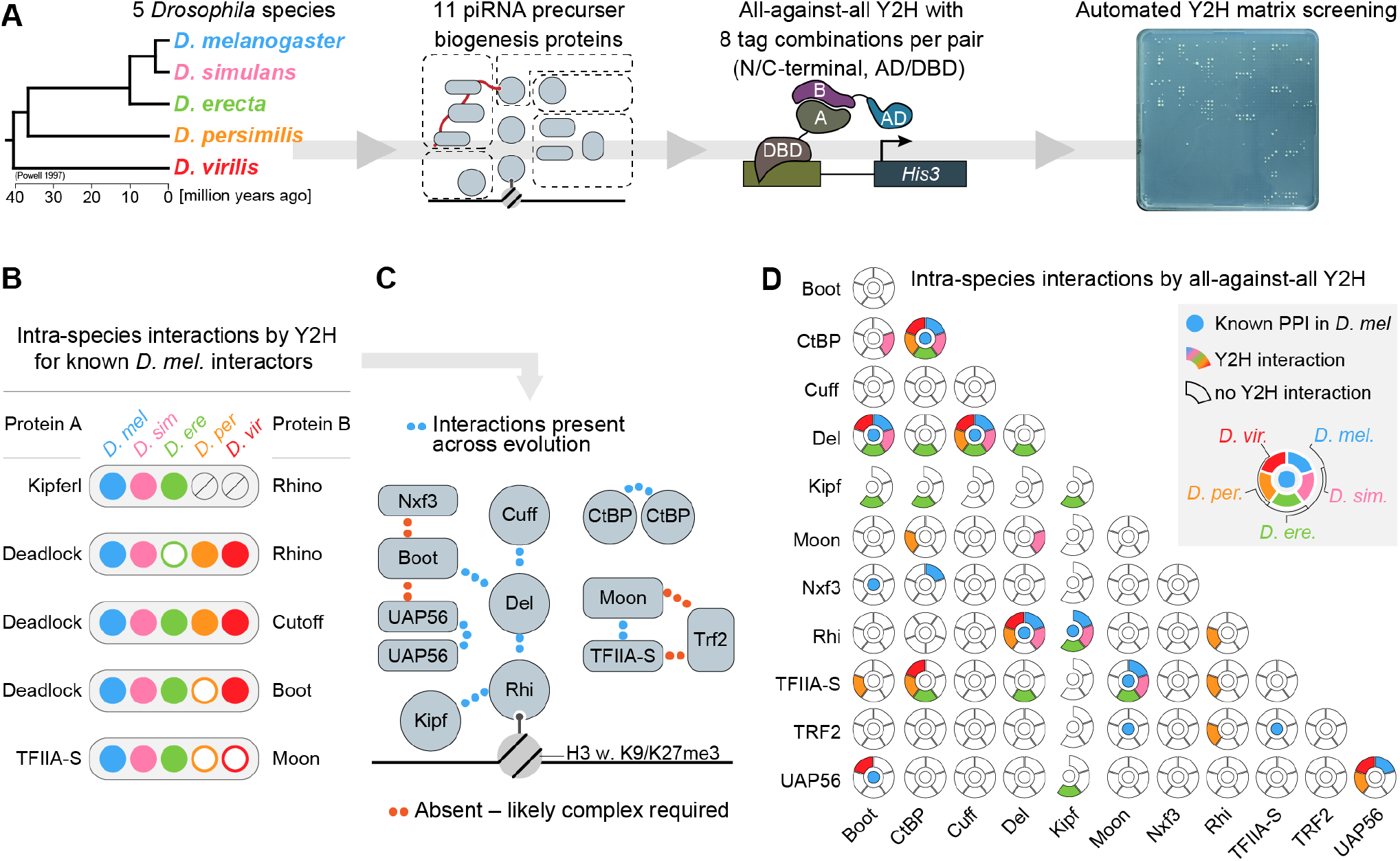
A yeast-two-hybrid screen uncovers the evolutionary biochemistry of piRNA biogenesis. (A) Schematic showing the experimental strategy of the yeast-two-hybrid screen. The major protein factors in germline piRNA precursor production are cloned from five *Drosophila* species spanning 40 million years of evolution and subsequently tested in all-against-all yeast-two-hybrid matrix screens. AD: activation domain. DBD: DNA-binding domain. See also Figure S2. The species color-code shown in the phylogenetic tree is the same throughout the paper. (B) Schematic summary of yeast-two hybrid results for interactions known from *D. melanogaster* amongst piRNA biogenesis factors within the same species (intra-species interactions). Full circles indicate detected interactions above the replication score threshold of 0.75, while empty circles indicate that interaction was not detected, or the score was below the threshold. Grey crossed-out circles for Kipferl denote absence of the gene in those species. (C) Schematic summary of the conserved interactions identified across the tested species phylogeny. (D) Schematic summary of yeast-two hybrid results for all tested intra-species interactions amongst piRNA biogenesis factors. Central blue dot: known interaction from *D. melanogaster*. Surrounding circle blocks indicate whether the interaction was detected (full block) or not (empty block). The color scheme follows the species annotation from (A).

Y2H experiments are prone to false negative results due to specific tagging, as fusions might interfere with folding or mask binding surfaces. To reduce the potential false negative rate^47,48^, we cloned all coding sequences as bait and prey, each fused C- and N-terminally with a DNA-binding (DBD) or a transcription activation domain (AD), respectively. Therefore, every protein is represented as four individual constructs, and every protein interaction is tested in eight combinations using different constructs^49^ **(Figure S2A-B)**. Bait proteins that did not show any interactions or, conversely, resulted in a set of well-sampled protein-protein interactions were screened two times. The other baits were screened three times, with three or four technical replicates each (**Figure S2C-F**). Excluding 10 auto-active bait and two auto-active prey constructs from the analyses, we examined a total of 89,088 individual pairs. For scoring Y2H interactions, we applied thresholds for colony size and technical as well as biological replication (**Figure S3**). Here, each unique configuration of a protein-protein interaction can lead to a maximum score of one. After combining all eight potential configurations for a protein pair, the sum of the scores can reach a maximum of eight. We then benchmarked the success of the screen by assessing protein-protein interactions previously described between piRNA precursor biogenesis factors in *D. melanogaster*. Using a conservative Y2H replication score cutoff of 0.75, we report 199 interacting protein pairs, 46 intra- and 153 inter-species interactions (**Figure S4**). The five most central protein interactions known from studies of the network in *D. melanogaster* were confirmed by Y2H (**Figure 2B**, blue dots). The four previously described interactions that were not detected by Y2H (Boot–Nxf3, Boot– UAP56, Trf2–Moon, and Trf2–TFIIA-S) (**Figure 2B-C**), are all part of multimeric complexes in the form of either Nxf3–Nxt1–Bootlegger–UAP56^35,36^ or Trf2–

TFIIA-S–Moonshiner^28^, thus likely preventing their detection via pairwise Y2H assays. Additionally, the previously described ability of CtBP and UAP56 to homodimerize^50,51^ was supported by Y2H signal in tests where the same protein was tagged as bait and prey (**Figure S4**). We therefore conclude that our Y2H screen recapitulated known pairwise interactions within the germline piRNA biogenesis pathway.

Not only are the vital *Drosophila* piRNA pathway genes evolving rapidly; some proteins found to be essential for the piRNA pathway in *D. melanogaster* are dispensable in closely related species^21^ revealing a remarkable evolutionary plasticity within the *Drosophila* piRNA pathway. We therefore investigated the degree of conservation of the core interaction network of germline piRNA biogenesis factors by comparing intra-species protein interactions across the investigated *Drosophila* species. Our Y2H assays showed that the central interactions connecting Deadlock to Rhino, Cutoff, and Bootlegger are detected in at least three of the four tested non-*melanogaster* species (**Figure 2B-C** and **Figure S4**). Similarly, we observed interaction between Rhino and Kipferl in all species harboring the *kipferl* gene. Furthermore, we did not find evidence for the proposed interaction between Cutoff and CtBP^29^ and only the *D. simulans* orthologs supported the proposed interaction between Moonshiner and Deadlock^28^ (**Figure 2B-D** and **Figure S4**). These results could reflect that (i) the Cutoff–CtBP and Moonshiner–Deadlock interactions are technically difficult to detect by Y2H, (ii) CtBP and Moonshiner are not involved in piRNA precursor biogenesis in some *Drosophila* species, or (iii) unknown and potentially evolutionarily labile interactions connect CtBP and Moonshiner to Rhino-dependent piRNA source loci in different *Drosophila* species. In sum, we uncovered an overall conserved interaction network structure amongst *Drosophila* germline piRNA biogenesis factors – with notable exceptions indicating evolutionarily labile recruitment of the CtBP and Moonshiner transcription factors.

### Cross-species analysis reveals diverse protein interaction evolution within the piRNA pathway

The conservation of a core protein interaction network within the piRNA pathway can be explained by at least two evolutionary trajectories: (i) protein interaction site conservation and (ii) protein interaction co-evolution, whereby compensatory mutations evolve that restore protein-protein interaction following sequence changes that otherwise diminish it. Interestingly, such interaction site co-evolution has previously been suggested for interactions between Rhino, Deadlock and Cutoff based on observed species incompatibilities between *D. melanogaster* and *D. simulans*^29,38^. We therefore looked for signatures of either interaction site conservation or co-evolution in our cross-species Y2H data representing evolutionary distances from 0 to 40 million years of divergence^52^ (**Figure 3A**). We reasoned that conserved interaction surfaces should yield Y2H interaction signals for proteins across the tested species (inter-species interactions), while co-evolving interactions should instead tend to show interaction signals between proteins from the same species (intra-species interactions) or from closely related species. We found that the Deadlock–Cutoff and Rhino–Kipferl interactions are detected at similarly high frequency in intra-species and inter-species interaction tests across all tested evolutionary distances (**Figure 3B** and **Figure S5A**), compatible with a model of conserved interaction surfaces. Conversely, the interactions of Deadlock–Rhino and Deadlock–Bootlegger showed a higher frequency of intra-species and short-distance interspecies interactions compared to tests between more distantly related orthologs (**Figure 3C-D**), indicative for a model of co-evolution. Consistent with these findings, co-evolution of the Deadlock–Rhino was previously proposed to have taken place based on comparative functional and structural studies across *D. melanogaster* and *D. simulans*^38,39^. Our Y2H data, however, also indicate functional innovation beyond the conserved interactions in the piRNA precursor biogenesis network, as evident from several species-restricted interactions supported by Y2H interactions detected only within one or two *Drosophila* species. For example, we found that *D. persimilis* Rhino interacts with the transcription factors TFIIA-S and Trf2 (**Figure S5B-C**). Such interaction with TFIIA-S and Trf2 from multiple species indicates that the molecular innovation enabling the interaction has taken place along the lineage leading to the *D. persimilis* Rhino. Similarly, we observed that Deadlock orthologs from *D. simulans* and *D. erecta* interact with Moonshiner from several species (**Figure 3E**). Finally, we also detected interaction between CtBP from most species and the Deadlock orthologs from *D. erecta* as well as *D. virilis* (**Figure 3F**). In sum, the presented Y2H data suggests that the individual protein interactions in the piRNA precursor biogenesis pathway follow separate evolutionary paths and are characterized by either conserved, co-evolving, or species-restricted interactions.

**Figure 3.**
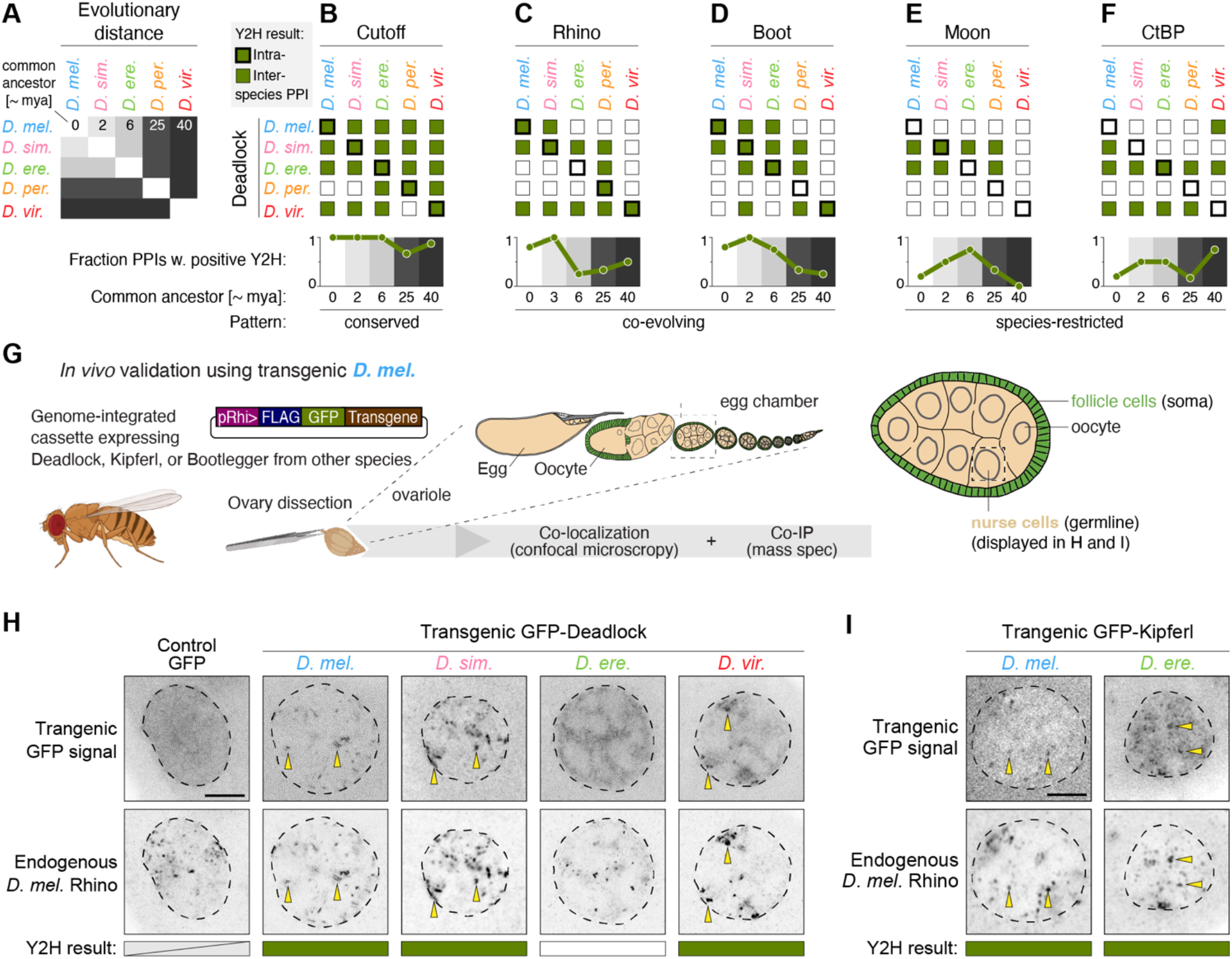
Protein-protein interaction evolution uncovered by cross-species interaction analysis. (A) Diagram indicating the estimated evolutionary distance between the tested Drosophila species. mya = million years ago. (B-F) Summary of interactions detected by yeast-two-hybrid between orthologs of Deadlock and Cutoff (B), Rhino (C), Bootlegger (D), Moonshiner (E), and CtBP (F). Intra-species (thick box outline) and inter-species (thin box outline) interactions are shown as filled green boxes, while empty boxes indicate absence of interaction detection above the replication score threshold. The plots below each summary matrix show the fraction of positive Y2H interactions split by the evolutionary distance between the source species. The fractions were calculated as the number of positive interactions divided by the number of tested interactions in each evolutionary distance group (see also A). Pattern: evolutionary signature of the interaction (see main text for details). (G) Schematic of the transgenic strategy testing interaction between proteins from different species (inter-species) *in vivo* by co-localization and co-immunoprecipitation analysis. (H-I) Confocal microscopy images showing the localization of endogenous *D. melanogaster* Rhino (anti-Rhi IF) and GFP-tagged transgenic Deadlock (H) or Kipferl (I) from the indicated species. Yellow arrows highlight co-localizing foci of endogenous Rhino IF signal and transgenic GFP-tagged proteins. Dashed line: nuclear border as determined by DAPI staining (see Figure S5D-E). Scale bars indicate 5 µm.

The protein-protein interactions that enable piRNA precursor production take place in *Drosophila* germline cells in a molecular environment very different from that of the yeast cells used in Y2H assays. When assayed by confocal microscopy, the investigated proteins accumulate in distinct foci within the ovarian germline nurse cell nuclei^23,24,27,28,33,35,36^. These foci represent major piRNA source loci and the accumulation of the individual proteins depends on the protein-protein interactions that directly or indirectly connect them to Rhino bound to the locus chromatin (**Figure 1A**). To probe how well our Y2H data translate to functionally relevant protein interactions *in vivo*, we generated transgenic *D. melanogaster* flies expressing GFP-tagged Bootlegger, Deadlock or Kipferl specifically in germline cells driven by the *rhino* promoter of *D. melanogaster*, which facilitates transgene expression similar to the endogenous *rhino* gene (ref. 27, **Figure 3G**). To assay protein interactions in their native environment, we first performed immunofluorescence confocal imaging of nurse cell germline nuclei and monitored the co-localization of the transgenic GFP-fusions with endogenous *D. melanogaster* Rhino. Consistent with the Y2H data, we observed that Deadlock from *D. melanogaster, D. simulans*, and *D. virilis*, but not from *D. erecta*, colocalizes with endogenous *D. melanogaster* Rhino (**Figure 3H** and **Figure S5D**). Of note, *D. simulans* Deadlock was also previously observed to co-localize with endogenous Rhino when expressed in *D. melanogaster*^38^. Similarly, we observed complete congruence of Y2H data and *in vivo* protein colocalization for the tested interactions of Rhino–Kipferl (**Figure 3I** and **Figure S5E**) and Deadlock–Bootlegger (**Figure S5F**, Bootlegger localization to Rhino foci relies on Deadlock). We conclude that patterns of conserved as well as co-evolving protein interactions inferred from our Y2H data are recapitulated by *in vivo* protein co-localization assays.

### Recurrent protein network rewiring identified by species-restricted protein interactions

In addition to the conserved or co-evolving interactions that place Deadlock and Bootlegger at central molecular hubs in piRNA precursor biogenesis (**Figure 2C**), our Y2H data also suggest that both proteins take part in several species-restricted protein-protein interactions. For example, *D. simulans* Bootlegger and *D. erecta* as well as *D. virilis* Deadlock showed highly reproducible interaction with CtBP in some species but not others (**Figure 2D, Figure 3F**, and **Figure S4**) and Deadlock from both *D. simulans* and *D. erecta* interacted with Moonshiner from several species (**Figure 3E** and **Figure S4**) but not others. To more directly characterize such species-restricted protein-protein interactions *in vivo*, we next performed co-immunoprecipitation coupled to mass spectrometry analysis (co-IP/MS) using ovary lysates from the transgenic *D. melanogaster* fly lines expressing Bootlegger and Deadlock proteins from various species. Co-IP/MS analysis of C-terminally tagged Bootlegger recapitulated interactions identified in previously published coIP/MS data, such as with Nxf3^36^ (**Figure S6A-B)**. The C-terminally tagged bootlegger also co-precipitated with all three PIWI-clade Argonaute proteins (Piwi, Aub, Ago3, **Figure S6A-B**), suggesting direct coupling of piRNA precursor export and cytoplasmic processing to mature piRNAs. Conversely, N-terminal tagging somehow disrupts the strong co-IP association of Bootlegger with the piRNA precursor export complex proteins UAP56, Nxt1, and Nxf3 (**Figure 4A-B** and **Figure S6A-B**). This disruption, however, did not perturb the species-restricted interactions detected by Y2H, as coIP/MS of N-terminally tagged *D. simulans* Bootlegger showed prominent co-precipitation of both Deadlock and CtBP (**Figure 4B, D**). In sum, we demonstrate the species-restricted robust interactions between *D. simulans* Bootlegger and Deadlock as well as CtBP by both Y2H and inter-species coIP/MS analysis from transgenic flies.

**Figure 4.**
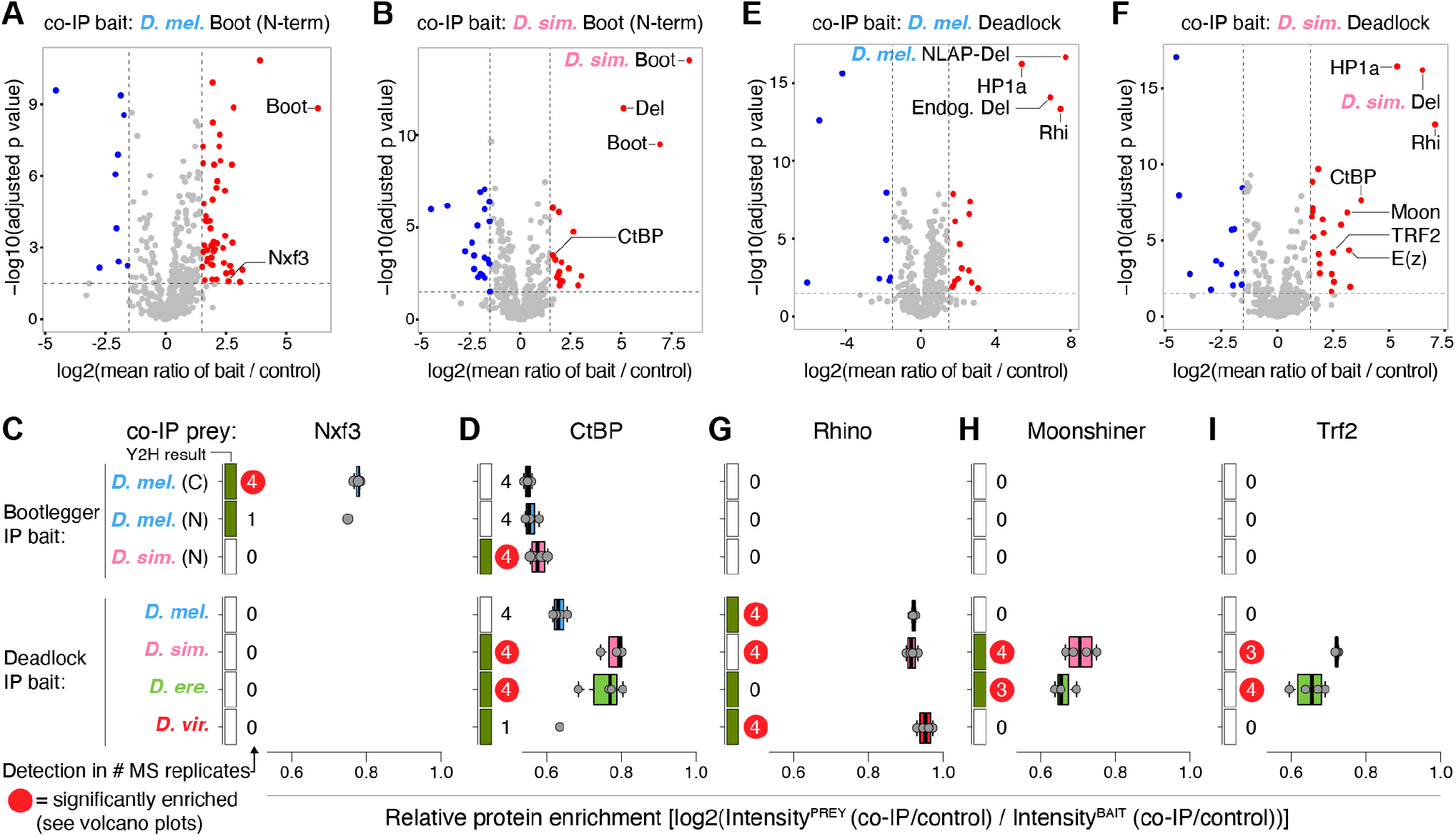
Cross-species *in vivo* Co-IP/MS shows innovation of protein interactions in piRNA biogenesis. (A-B) Volcano plots displaying co-IP/mass spectrometry data from the co-IP baits indicated above each plot. The x axes show log2 fold change between bait and control IP averaged over four biological replicates with each two technical replicates. The y-axis display the negative log10 of adjusted p values from t-tests for enrichment in bait compared to control co-IP. (C-D) Box plots showing co-IP protein binding enrichment relative to GFP-tagged bait binding for the indicated bait and prey proteins. Rectangles at the left y-axis indicate whether the protein interaction was detected by yeast-two-hybrid (filled green) or not (empty white). Numbers denote the number of co-IP biological replicates in which the prey protein was detected. Red circles indicate that the prey protein was determined to be statistically significantly enriched (see A-b, E-F and Figure S6B-D). Grey dots represent enrichment values for individual biological replicates averaged over two technical replicates. (E-F) Volcano plots similar to (A-B). (G-I) Box plots similar to (C-D).

In the *D. melanogaster* Deadlock coIP/MS analysis, Rhino, and HP1a were co-immunoprecipitated (**Figure 4E**), whereas Bootlegger and Cutoff were not, despite the strongly supported interaction between these proteins based on Y2H (**Figure 3B,D** and **Figure S4**). This suggests that tagged Deadlock transgenes may only partially recapitulate endogenous Deadlock interactions. However, consistent with our co-localization analysis (**Figure 3H**) and the proposed co-evolution model, Rhino protein was identified in coIP/MS analysis of *D. simulans* and *D. virilis* Deadlock, but not in *D. erecta* Deadlock coIP/MS (**Figure 4E-G** and **Figure S6C-D**). In addition, the species-restricted robust interaction between Deadlock and Moonshiner identified by Y2H (**Figure 3E** and **Figure S4**) was supported by pronounced co-precipitation of Moonshiner as well as the Moonshiner binding partner Trf2 (**Figure 4F,H-I** and **Figure S6C**). Given the Y2H-based *D. persimilis*-specific interactions between Rhino and TFIIA-S as well as Trf2 (**Figure S4**), our data collectively indicate recurrent innovation of the protein-protein interactions leading to RNA Polymerase II recruitment at germline piRNA source loci (see Discussion).

Surprisingly, we also observed notable co-precipitation of Enhancer of Zeste (E(z)) protein in coIP/MS analysis of *D. simulans, D. erecta* and *D. virilis* Deadlock (**Figure 4F** and **Figure S6A**,**C-D**). E(z) is the catalytic subunit of the Polycomb Repressive Complex 2 (PRC2), which deposits the repressive H3K27me3 mark^53–55^. Of note, amongst the seven known PRC2 proteins, only E(z) was observed to be enriched in the Deadlock coIPs (**Figure S6A**), indicating that E(z) may be recruited to Deadlock alone. H3K27me3 marks were recently shown to mediate Rhino binding at many germline piRNA source loci in *Drosophila*^26^. The identified connection between Deadlock and E(z) therefore has potential implications for piRNA source locus regulation, as it implies that H3K27me3 may be deposited as a consequence of Rhino–Deadlock association with loci rather than acting upstream of such binding (see Discussion).

Finally, coIP/MS analysis uncovered co-precipitation of *D. melanogaster* CtBP with Deadlock from *D. simulans* and *D. erecta*, and – substantially weaker – with Deadlock from *D. melanogaster* but not with Deadlock from *D. virilis* (**Figure 4D-F** and **Figure S6C-D**). Our Y2H data indeed support direct interaction between CtBP and both *D. simulans* and *D. erecta* Deadlock (**Figure S4**). Furthermore, *D. virilis* Deadlock interaction with CtBP was strongly supported by Y2H data (**Figure S4**), but was not recapitulated by coIP/MS (**Figure 4D** and **Figure S6A**,**D**). Notably, the co-precipitations of CtBP with *D. simulans* and *D. erecta* Deadlock were also accompanied by robust interaction with the Moonshiner–TFIIA-S–Trf2 complex (**Figure 4H-I**). Across all tested species, low but consistent Y2H scores supported interaction between TFIIA-S and CtBP (**Figure S4**). We therefore speculate that robust Deadlock-Moonshiner interaction may stabilize CtBP interactions with Deadlock through additional CtBP-TFIIA-S interactions.

In sum, our Y2H data together with *in vivo* colocalization and coIP/MS analyses reveal multiple species-restricted protein interactions amongst the proteins facilitating transcriptional regulation at germline piRNA source loci, implicating rapid functional innovation of the responsible protein-protein interaction surfaces.

### Short linear motif evolution facilitates mechanistic divergence in CtBP recruitment

To understand which changes in protein sequence may facilitate the observed innovation in protein-protein interactions, we next focused on the rewiring of CtBP interactions. Close inspection of CtBP Y2H interaction data revealed a highly variable pattern of interaction partners across the investigated species (**Figure 5A-G**). CtBP dimerization, which is known to facilitate its repressive functions^50^, was robustly detected within and between all species (**Figure 5A** and **Figure S4**); however, interactions with TFIIA-S, Deadlock, Nxf3, Kipferl, Moonshiner, and Bootlegger were found in only few or a single species (**Figure 5B-G** and **Figure S4**). The CtBP interaction with TFIIA-S was found across all species but *D. melanogaster*, while only *D. erecta* and *D. virilis* Deadlock interacted with the CtBPs. NXF3, Kipferl Moonshiner and Bootlegger interaction with CtBPs was specific for *D. melanogaster, erecta, persimilis*, and *simulans*, respectively (**Figure 5 D-G**)

**Figure 5.**
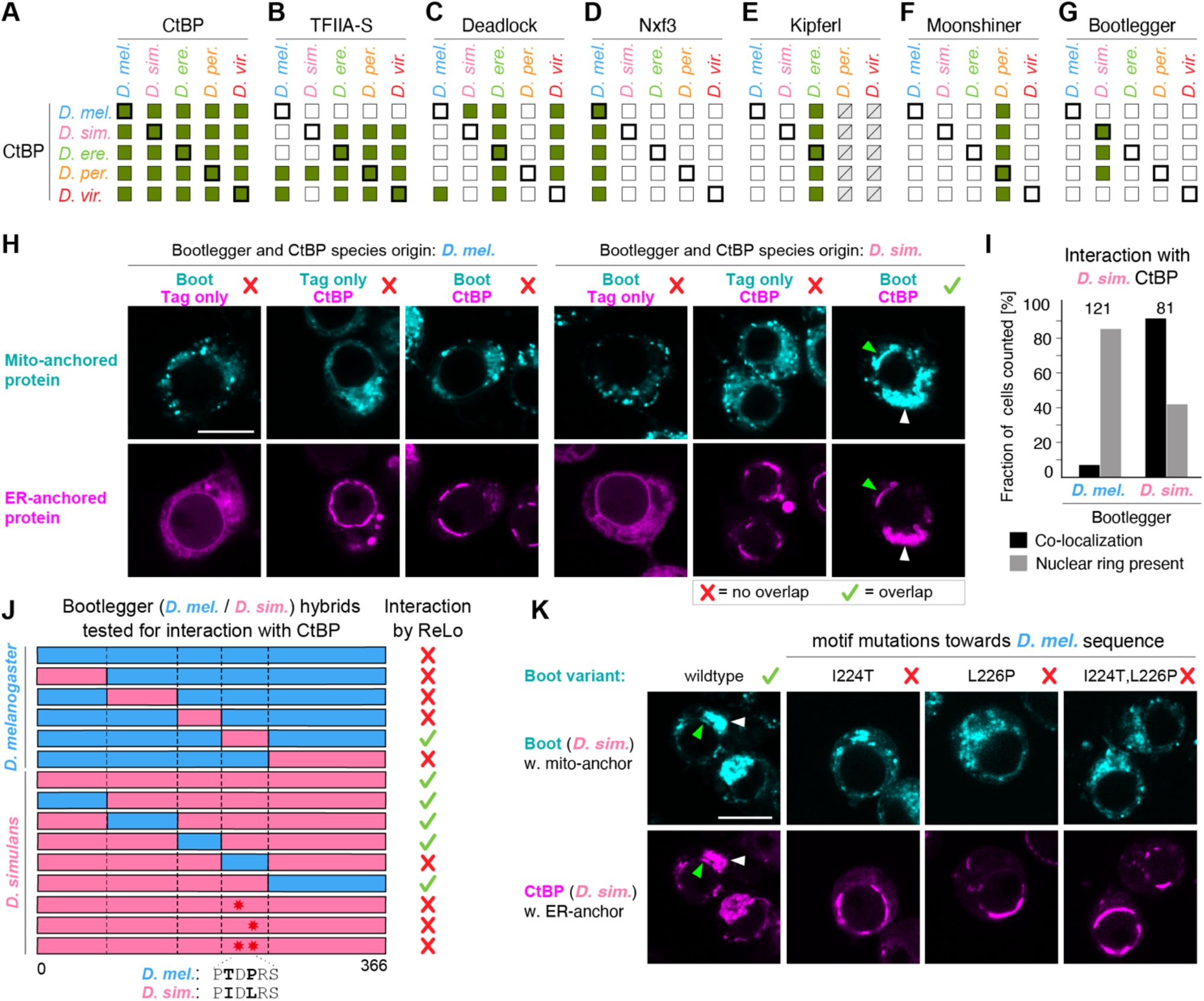
Recurrent innovation of CtBP interaction with piRNA biogenesis factors. (A-G) Summary of interactions detected by yeast-two-hybrid between orthologs of CtBP and CtBP (A), TFIIA-S (B), Deadlock (C), Nxf3 (D), Bootlegger (E), Kipferl (F) and Moonshiner (G). Intra-species (thick box outline) and inter-species (thin box outline) interactions are shown as filled green boxes, while empty boxes indicate absence of interaction detection above the replication score threshold. (H) Fluorescence microscopy images of ReLo protein interaction assays testing interaction between CtBP and Bootlegger proteins from *D. melanogaster* (left panel) and *D. simulans* (right panel). Shown are representative cells from >10 images. Green arrows indicate protein accumulation in interrupted ring structures around the nucleus. White arrows show protein accumulation at cytoplasmic sites. Scale bars indicate 8 µm in size. Red cross and green check mark denote whether protein interaction was concluded or not, respectively, based on the ReLo assays. (I) Bar plot displaying the quantification of ReLo co-localization and nuclear ring-like phenotypes in 202 individually evaluated cells. (J) Schematic representation of domain-swap constructs between the *D. melanogaster* (blue) and *D. simulans* (pink) Bootlegger orthologs with the interaction results indicated to right by red crosses or green check marks. Black stars indicate the tested interaction site point mutants. (K) Fluorescence microscopy images of ReLo protein interaction assays testing point mutants of *D. simulans* Bootlegger for interaction with CtBP. Displayed as for (H).

Importantly, given the high conservation of the CtBP amino acid sequence across the tested species (**Figure S7**), Y2H assays with CtBP from each species serves as further replicates, attesting to the robustness of the identified interaction signals.

To understand the molecular mechanism underlying recurrent CtBP interaction rewiring, we decided to characterize the interaction between *D. simulans* Bootlegger and CtBP (**Figure 4B** and **Figure 5G**). To this end, we used a qualitative, fluorescence-based re-localization protein-protein interaction assay (ReLo)^56–58^. Here, plasmids encoding candidate interacting proteins fused to a fluorescent protein and a membrane associated anchor are co-transfected into *Drosophila* S2R+ cells. Protein-protein interactions are then assayed by colocalization analysis using live cell confocal imaging. To adapt the ReLo assay to nuclear proteins, we fused the proteins with either a mitochondrial outer membrane anchor (Daed^TMM^) or an endoplasmic reticulum (ER) membrane anchor (OST4), thus forcing the fusion proteins to remain in the cytoplasm. Confirming our results from Y2H and coIP/MS assays, we observed that Bootlegger-mCherry-Daed^TMM^ displays co-localization with ER-anchored OST4-GFP-CtBP protein when the assay was done using *D. simulans* ortholog, while this was not observed for the respective proteins from *D. melanogaster* (**Figure 5H-I**), thus further support the presence of a species-restricted interaction between Bootlegger and CtBP in *D. simulans*. Of note, while control OST4-GFP localized to cytoplasmic endoplasmic reticulum-like regions, OST4-GFP-CtBP protein accumulated in ring structures around the nucleus with some cytoplasmic foci (**Figure 5H-I**). We speculate that such accumulation of the CtBP fusion protein reflects a strong nuclear localization signal, which traps the proteins at the nuclear envelope due to its simultaneous ER-association.

To identify the *D. simulans* Bootlegger interaction surface to CtBP, we generated a series of chimeric expression constructs with systematic domain swaps between *D. melanogaster* and *D. simulans* Bootlegger (**Figure 5J** and **Figure S8A**). ReLo assays using these constructs revealed that a central 50-amino acid region in *D. simulans* Bootlegger is necessary and sufficient for interaction with CtBP (**Figure S8A-C**). Inspection of this region uncovered a putative PIDLRS short linear motif (**Figure S8D**), previously identified to facilitate interaction between human hCTBP2 protein and the HCP2 polycomb group developmental regulator^59^. To test whether the PIDLRS motif is required for *D. simulans* Bootlegger interaction with CtBP, we mutated either one or both of the amino acids in the motif that differ from the corresponding *D. melanogaster* sequence (I224T, L226P). ReLo assays using *D. simulans* Bootlegger with single or double *D. melanogaster*-like motif mutation showed complete loss of co-localization (**Figure 5J-K**), suggesting that a short linear binding motif specific to *D. simulans* Bootlegger is necessary and sufficient for binding to CtBP. Notably, in *D. erecta* Deadlock, for which both Y2H and coIP/MS data strongly supported robust interaction with CtBP (**Figure 3F** and **Figure 4B**), we also identified a variant CtBP interaction motif (PLDLS) (**Figure S8E**), known to facilitate CtBP interaction in animals^60^. We conclude that recurrent evolution of short linear binding motifs contributes to rapid innovation of the piRNA pathway protein-protein interaction network in *Drosophila*.

## DISCUSSION

Here, we investigated the biochemical consequences of rapid evolution within 11 piRNA precursor biogenesis genes across five *Drosophila* species spanning an evolutionary distance of 40 million years. Using a well proven, high quality Y2H matrix array screening setup, we comprehensively tested the physical protein interactions of the 11 piRNA pathway members of *D. melanogaster, D. simulans, D. erecta, D. virilis* and *D. persimilis* in a pairwise, all-versus-all screen, yielding both comprehensive within (intra) and cross-species (inter) protein interaction data. Based on a high-quality data set of 199 protein interactions (46 intra and 153 inter-species), we elucidated evolutionary patterns of interaction, including strict conservation, coevolution, and species-restricted interactions. These inferences were further validated and characterized using *in vivo* co-immunoprecipitation and cell culture assays.

### Innovation of protein-protein interactions underlie rapid evolution in the piRNA pathway

Our work links adaptive evolution of protein sequence and the underlying innovations in biochemical functionality. According to evolutionary theory, essential cellular pathways should undergo strong negative (purifying) selection because changes in proteins could lead to inviability. However, evolutionary studies have found that genes supporting certain functions, such as immune response^61–64^ and defense against viruses and transposons^14,19,65^ often evolve rapidly with selection *for* amino acid-changing variants (positive selection)^66,67^. Yet, the innovation in biochemical functionality that takes place in such rapidly evolving, albeit essential genes has been characterized in only a few cases, such as the primate retroviral restriction factor TRIM5α^68^, mouse coat color adaptation^69^, and toxin resistance evolution in insects^70^. Furthermore, proteins often function in complex pathways and protein-protein interaction networks, and it remains unclear how adaptive evolution shapes functional pathways composed of multiple connected proteins.

We also find that the adaptively evolving genes supporting *Drosophila* germline piRNA biogenesis, in addition to several conserved protein interactions, display at least two layers of innovation of biochemical functionality. First, we showed that while most interactions known from *Drosophila melanogaster* are conserved in the other *Drosophila* species, several show evidence of protein complex interface co-evolution (**Figure 6A**). Second, we observed rewiring affecting protein interactions in the recruitment of CtBP and Moonshiner–TFIIA-S–Trf2 transcriptional regulators. In this way, the recruitment of these effectors via protein-protein interaction is conserved, but the identity of the connected proteins is evolutionarily fluid.

**Figure 6.**
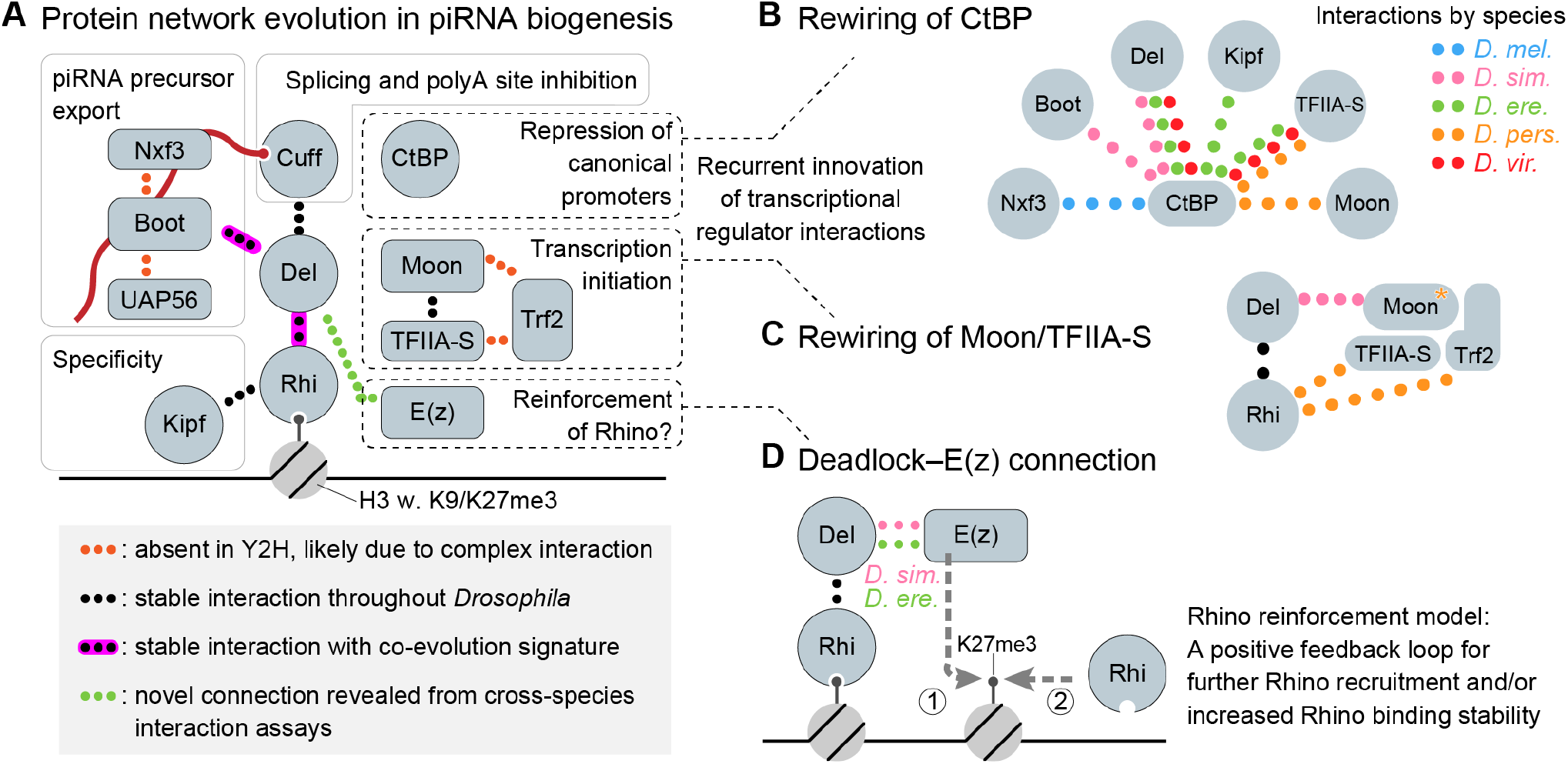
Evolutionary biochemistry of the germline piRNA biogenesis factor network. (A) Schematic model summarizing the identified evolutionary characteristics for individual protein-protein interactions within the germline piRNA precursor biogenesis network. Dotted lines represent detected protein-protein interactions. See legend for details. (B-C) Schematic models of the observed evolutionary innovation of protein interactions between core piRNA biogenesis factors and the CtBP (B) and Moonshiner–TFIIA-S–Trf2 (C) transcriptional regulators. Colors of the dotted lines indicate within which species the protein interactions were detected. (D) Proposed model for the functional role of the observed interaction between E(z) and Deadlock from *D. simulans* and *D. erecta*. We speculated the E(z) recruited via Deadlock to Rhino-bound H3K9me3 histones may reinforce Rhino binding through increased Rhino recruitment and/or increased residence time (see also Discussion).

Of note, there are also rapidly evolving proteins in the network for which we did not uncover innovation in protein-protein interactions (Cutoff and Kipferl). This observation reflects three limitations of the current study: (i) innovation could have occurred for intermolecular interactions beyond the investigated network, (ii) the underlying innovation may concern biochemical properties not tested in the present study, such as protein folding, stability, or fine-tuning of protein-protein interaction affinity, which is not well-captured by the more qualitative nature of Y2H assays^71^, and (iii) it is not possible to directly link the detected positive selection signatures to specific innovations in the interaction network. Future research using unbiased interactome profiling, quantitative affinity measurements, and broad species sampling to increase temporal resolution holds the promise to address these limitations and the connected open questions.

### Species-restricted recruitment of transcriptional regulators to piRNA source loci

Our Y2H data uncovered multiple species-restricted interactions between CtBP and different proteins in the Rhino interaction network (Nxf3, Deadlock, Bootlegger, Kipferl, TFIIA-S, and Moonshiner) (**Figure 6A**). We interrogated the *in vivo* relevance of these interactions by co-IP/MS of *D. simulans* Bootlegger and Deadlock from *D. simulans* and *D. erecta* and revealed strong congruence between the Y2H and coIP/MS data (**Figure 4**). Furthermore, we were able to identify a short linear interaction motif specific to *D. simulans* Bootlegger that is necessary and sufficient for interaction with CtBP (**Figure 5 and Figure S8**). A similar motif has also evolved in *D. erecta* Deadlock (**Figure S8**), which we by both Y2H and coIP/MS found to interact with CtBP. Together, our results reveal extensive evolutionary innovation in the recruitment path (i.e. interaction rewiring) of CtBP to Rhino-bound piRNA source loci (**Figure 6B**). CtBP recruitment to piRNA source loci has been shown to silence the activity of canonical DNA-encoded promoters, while not affecting the non-canonical heterochromatin-dependent transcription initiated via Moonshiner^29^.

In addition to rewiring of CtBP recruitment, we also identified species-restricted interactions connecting the Moonshiner–TFIIA-S–Trf2 complex to the Rhino interaction network (**Figure 6C**). While it remains unclear if the previously proposed recruitment of Moonshiner via Deadlock in *D. melanogaster*^28^ is supported by direct interaction between the two proteins, our Y2H as well as coIP/MS results show a clear interaction between Moonshiner and Deadlock from *D. simulans* and from *D. erecta* (**Figure 4** and **Figure S6**). We propose that Moonshiner complex recruitment via direct interaction with Deadlock is central to the *Drosophila* piRNA pathway but that yet unknown functional variation between the investigated orthologs contributes to the observed differences in Deadlock– Moonshiner interaction. Furthermore, in *D. persimilis*, interactions were observed by Y2H between Rhino and both TFIIA-S and Trf2, suggesting that recruitment of the Moonshiner– TFIIA-S–Trf2 complex may be supported by interaction with Rhino in this species. Together with the rewiring of CtBP recruitment, these findings reveal that the interactions mediating recruitment of the key transcriptional regulators of germline piRNA source loci undergo recurrent evolutionary innovation. Interestingly, expression of *D. simulans* Cutoff in *D. melanogaster* was previously found to lead to increased association of Trf2 and and CtBP with Rhino-bound loci, suggesting that adaptation of Cutoff also contributes to the recurrent innovation of transcription factor recruitment^29^. Given that we did not detect direct interactions between Cutoff and such transcription factors, it seems likely that *D. simulans* Cutoff stimulates transcription factors recruitment indirectly or through complex formation with e.g. Deadlock. We propose that rewiring of CtBP and Moonshiner–TFIIA-S–Trf2 complex recruitment may reflect adaptive evolution to ensure transposon promoter silencing and content balancing of the piRNA pool during host–transposon arms race evolution (further discussed below).

While our well controlled Y2H matrix assay by experimental design defines the set of investigated protein-protein interactions, the coIP/MS experiments also allowed identification of novel protein interactions. These data uncovered an unexpected enrichment of the E(z) protein in co-immunoprecipitations of Deadlock from *D. simulans, D. erecta*, and *D. virilis*, which we did not find with *D. melanogaster* Deadlock (**Figure 4** and **Figure S6**). E(z) is the catalytic subunit of the Polycomb Repressive Complex 2 (PRC2) complex and is responsible for the methylation of lysine 27 on the N-terminal tail of Histone 3 (refs. 72,73). H3K27me3 modification by PRC2 was recently found to co-occur with H3K9me3 modifications at Rhino-bound piRNA source loci^26^, which together was proposed to define a dual histone code that specifies Rhino binding at a subset of piRNA source loci. Based on our observed connection between Deadlock and E(z) in several *Drosophila* species, we speculate that E(z) recruitment – potentially outside the context of a PRC2 complex – may function in a Rhino-binding reinforcement loop (**Figure 6D**). Specifically, initial recruitment of Rhino may lead to Deadlock-mediated E(z) recruitment and subsequent establishment of H3K27me3 marks, which then increases Rhino recruitment or binding stability^26^ to stabilize the genomic region as a piRNA source locus.

### Possible genetic drivers of adaptive evolution and biochemical innovation in the piRNA pathway

Our work uncovers biochemical innovations within an adaptively evolving protein network at an unprecedented scale. What might be the biological advantage of such rapid biochemical innovation in a network of essential genes? At least three broader evolutionary forces may contribute to such rapid evolution despite the essential function of the piRNA pathway. First, the adaptive walk model of evolution^74,75^, states that young genes are expected to evolve faster and with larger mutational fitness effect because they are further from their fitness optimum compared to older genes. Adaptive walk effects may therefore contribute to the rapid evolution of young, Drosophila-restricted piRNA pathway genes such as Rhino, Deadlock, Cutoff, Moonshiner, Bootlegger and Nxf3. Second, the piRNA autoimmunity model^22,76–79^ proposes that the rapid evolution of the genomic loci giving rise to piRNAs would frequently give rise to piRNAs that target important host genes for silencing. Under this model, such self-targeting would drive adaptation for changes in piRNA pathway proteins that reduce the host gene silencing, e.g. by altering the piRNA biogenesis mechanism or the silencing potency. Third, the evolutionary arms race model suggests that recurrent rapid evolution of piRNA pathway genes occurs in response to transposon evolution towards evasion of piRNA silencing or ‘anti-silencing’ by inhibition of the piRNA pathway. While such “Red Queen” evolution^80^ has been proposed based on observations of functional diversification of piRNA pathway genes between closely related species^29,38,81^, only the recent identification of a *Drosophila* transposon expressing piRNAs that target a piRNA pathway factor for silencing^82^ supports the existence of such piRNA pathway inhibitors. Finally, our data also suggest a fourth, ‘piRNA locus balancing’, model to explain rapid piRNA pathway evolution. This model rests on the observation that germline piRNA source loci differ in their dependence on specific piRNA precursor biogenesis factors. Some source loci, for example, are expressed independently of Moonshiner and Trf2 and instead rely on DNA-encoded promoters^28^. Other loci do not require the zinc-finger protein Kipferl^27^ and a few germline piRNA source loci (e.g. *cluster20A*) are expressed independently of any of the factors in the Rhino interaction network^23,24,27,28,36^. We propose that altering the interactions between proteins at germline piRNA clusters could impact piRNA production from specific loci. Depending on current transposon activity in a population, altered piRNA sub-pools could provide a mechanism to shift the target focus between transposon families. This model is thus consistent with the recent observation that while piRNA clusters tend to be syntenic between *Drosophila* species, they share little sequence content and instead contain large insertions of recently active transposons^83^.

In sum, our study of the evolution of protein-protein interactions across *Drosophila* evolution has uncovered principles of interaction evolution within an adaptively evolving protein network. Our findings uncover how pathway function is maintained despite rapid biochemical innovation and lay the foundation for better mechanistic understanding of the biochemically labile piRNA pathway and the evolutionary forces driving such innovation. Future studies linking molecular mechanistic characterization and evolutionary analyses will be key to uncover which evolutionary forces are driving the rapid evolution and biochemical innovation of the piRNA pathway in *Drosophila*. Amongst such studies, further probing for transposon-encoded silencing evasion or anti-silencing mechanism as well as comparative investigations of piRNA source locus regulation in different *Drosophila* species promise to be revealing.

## Supporting information

Table_S1_PAML

Table_S2_Protein_IDS_Y2H

Table_S3_Flies_used_in_this_study

## ACKNOWLEDGEMENTS

We much appreciate the helpful comments on earlier versions of the manuscript from Julius Brennecke and Anne F. Nielsen. We thank the Nikon Imaging Center of the University of Heidelberg for access to microscopes. We thank Jutta Metz for generating the pAc5.1-mCherry-MLS vector. S.R. was supported by an EMBO scientific exchange grant (#10132) and the Aarhus University GSNS Mobility Grant. Work in the Jeske lab was funded by the Emmy Noether Program of the German Research Foundation (DFG; JE-827/1-1 to M.J.). The Andersen group is supported by the Novo Nordisk Foundation (NNF18OC0030954), the Independent Research Foundation Denmark (9064-00056B), and an AIAS Marie Curie CO-FUND Fellowship (P.A). The work was also supported by BioTechMed-Graz, project DYNIMO, and the Austrian Science Fund, FWF (doi: 10.55776/P30162, doi: 10.55776/P34316), as well as the Field of Excellence BioHealth - University of Graz.

## AUTHOR CONTRIBUTIONS

S.R. led the project, performed the experimental work, analyzed data except from the contributions listed below, and drafted the manuscript.

S.M. performed mass spectrometry sample preparation after co-IPs and data processing.

S.F. contributed to Y2H screen preparation. S-Y.L. performed the evolutionary analyses.

M.A. contributed to plasmid cloning and validation.

H.K.S. supervised and contributed to the ReLo assay data collection.

A.E. performed fly genetics and contributed to ovary collection for co-IPs

M.J. contributed to ReLo assay data analysis and manuscript editing and supervised H.K.S.

M.T.L. contributed to evolutionary data analysis and manuscript editing and supervised S-Y.L.

U.S. contributed to Y2H and mass spectrometry data analysis as well as manuscript editing and supervised

S.M. and S.F.

P.A. conceived the study, contributed to data analysis, edited the manuscript, and supervised S.R. A.E, and M.A.

## MATERIAL AND METHODS

### Evolutionary analyses of germline piRNA cluster transcription and export genes

dN/dS between *D. melanogaster* and *D. simulans* was calculated using the software, DnaSP (v6; ref. 84). The *D. melanogaster* alleles and the *D. simulans* alleles were obtained from the genome annotations dmel r6.53 on FlyBase and the Prin_Dsim_3.1 on NCBI, respectively. To test for positive selection, a phylogeny-based test using the codeml program in PAML (v4; ref. 85) was implemented. The likelihood of models M7 (neutral: the dN/dS values fit a beta distribution between 0 and 1) and M8 (non-neutral: M7 parameters plus dN/dS > 1) were compared, using the F3×4 model of codon frequency. The degrees of freedom for each test were set to 2 and the log likelihood ratio test was inferred as significant if the p-value of the Chi-squared test was less than 0.05. All orthologs were extracted from either FlyBase or NCBI. PAML results and species are listed in **Figure S1**.

### Plasmid cloning by Gibson assembly

Plasmids for ReLo assays and fly injections were assembled, designed in SnapGene (v7.0.3), and cloned via Gibson reactions (NEB; E2611L) according to the instruction manual with 20 - 25 nt of fragment overhang by default. Reactions were pipetted on room temperature, immediately shifted to 50°C after addition of the reaction mix for 60 minutes and afterwards transformed into DH5α chemically competent bacteria. 50 µL bacteria were thawed on ice, 2 µL of the pre-chilled Gibson reaction was used to transform bacteria. The mixture was gently flicked and then incubated for 20 minutes at room temperature, transferred to 42°C for 42 seconds and then chilled on ice for 2 minutes. 200 µL of standard LB medium (10 g/L NaCl, 10 g/L peptone from casein, 5 g/L yeast extract) was added and the cells were incubated at 37°C at 550 rpm for 1 hour. Cells were then plated on Antibiotic containing LB-agar (LB medium with 15 g/L agar) plates and incubated over night at 37°C.

### Plasmid cloning for yeast-two-hybrid assays

Orthologous genes to the D. melanogaster piRNA pathway genes were identified as reciprocal best hits from BLAST^86^ searches. Of note, we did not find evidence of synteny at the reciprocal best BLAST hits for *D. melanogaster* Moon (DmMoon) and Nxf3 (DmNxf3) in *D. persimilis*, indicating potential relocation by recombination or duplication and subsequent loss of the original copy. Coding sequences of the orthologs were pulled from NCBI (see *Coding sequences used for the Y2H screen*) and synthesized by Twist Bioscience, with exception of all Deadlock orthologs which were amplified from cDNA of *Drosophila* ovaries. The 53 coding sequences of five different *Drosophila* species were cloned into pDONR221 (gift of Will Garland, Torben Heick Jensen lab, Aarhus University) with BP clonase II (Thermo Fisher, #10348102). Each CDS was then cloned into two Gateway compatible bait (Addgene #111232 & #111236) and two prey (Addgene #111233 & #111237) vectors^49^ with LR clonase II (Thermo Fisher; 11791100). LR reactions were performed according to standard protocols and 2 clones of each construct were picked for DNA mini preparation (Refs. 87,88). Each plasmid was digested with Bsp1407l to assess expected gene size by agarose electrophoresis.

### Coding sequences used for the Y2H screen

All Del sequences were amplified from cDNA of the respective species. Female flies were cultivated as described in *<u>Co-IPs from FLAG-tagged proteins for mass spectrometry</u>* and ovaries were lysed in Trizol reagent (Fisher Scientific, 15596018). Total RNA was isolated using Trizol reagent following the manufacturer’s instructions. 1 µg of total RNA was DNase treated and reverse transcribed using the iScript gDNA clear cDNA Synthesis kit (BioRad, 1725035). CDSs were then amplified with sequence specific primers annealing at the start (forward) and before the stop codon (reverse), containing overhangs for Gibson assembly. 500 ng of reverse transcribed cDNA was used as input. Correctly sized amplicons were used for Gibson reactions to clone them into Gateway vectors (see *<u>Plasmid cloning for yeast-two-hybrid assays</u>*). All other proteins were ordered as gene fragments (< 1800 bp) or clonal genes (> 1800 bp) from Twist Bioscience, CA, USA.

### Yeast growth and transformation

Yeast strains were grown on either yeast-extract-peptone-dextrose-adenine (YPDA) medium (content (w/v): 1% bacto-yeast extract; 2% bacto-peptone; 2% glucose; 0.004% Adenine) or supplemented nitrogen base (NB) (1.33% yeast nitrogen base (w/v)) for growth under selection. Supplemented NB was in each case added 2% Glucose; 0.004% Adenine; 0.002% Uracil and furthermore the following specific to the use case: (i) Growth of bait vectors: 0.002% each of Leucine, Histidine, and Methionine. (ii) Growth of prey vectors: 0.002% each of Tryptophane, Histidine, and Methionine. (iii) Mating Control: 0.002% each of Histidine and Methionine. (iv) Auto activity test of baits: 0.002% each of Leucine and Methionine.

A correctly sized replicate plasmid of the shuffled plasmids was used to transform Mat-a (bait vectors, Trp selection) or Mat-α (prey vectors, Leu selection). The bait strain genotype was *his3Δ300; trp1-901; leu2-3,112; ade2; GAL4; can1; cyh2; URA3::(lexAop)8-*GAL1TATA*-lacZ; LYS2::(lexAop)4-*HIS3TATA*-HIS3*. The prey strain genotype was *his3Δ300; trp1-901; leu2-3, 112; ade2; lys2-801am; gal4; gal80; cyh2; can1; ura3::(lexAop)8-*GAL1TATA*-lacZ; LYS2::(lexAop)4-*HIS3TATA*-HIS3; ADE2::(lexAop)8-*GAL1TATA*-URA3*.

Pre-transformation yeast cultures were grown over night at 30°C in 10 mL YPDA medium while shaking at 130 rpm. On the next day the ODs of yeast strains were measured, and 60 mL (60 mL are sufficient for one 96-well plate) main cultures were infected to OD 0.13 and grown as before. At OD 0,5 – 0,8 cells are harvested by centrifuging for 5 minutes at 1.000 g and washed once in PBS. In the meantime, flat bottom 96-well plates were prepared with ∼350 ng plasmid DNA, 25 µg salmon ssDNA (Sigma Aldrich, D1626). Yeast cells were resuspended in 2,2 mL Mix 1 (0.1 M Lithium Acetate; 5 mM Tris-HCl pH 8.0, 0.5 mM EDTA; 1 M Sorbitol) and incubated for 20 minutes at room temperature. Yeasts were resuspended again and 22 µL of the mix was added to each well, prepared with DNA. The mix was mixed via vortexing the plate. Next, 120 µL of Mix 2 (0.1 M Lithium Acetate; 10 mM Tris-HCl pH 8.0; 40% PEG3350) were added, wells were mixed, and the plates were incubated at 30°C for 30’. Then 16 µL DMSO were added, wells were mixed by vortexing, and plates were heat shocked at 42°C for 15 minutes. Finally, each transformant was stamped out four times. Four biological replicates of each transformed strain were maintained throughout the screen.

### Yeast-two-hybrid matrix screening

The basic workflow of the screen was performed as described^87,88^. All tables including raw counts and processing are available as Microsoft Access or Excel files on request. A custom gridding robot (kbiosystems) was used to stamp yeast in 96- and 384-well formats onto agar growth plates. Mating of haploid yeasts was performed on YPDA medium. For selective growth a nitrogen base medium, supplemented with amino acids and nucleotides was used. All solid trays and plates for yeast growth were supplemented with 2% Agar to solidify. Individual media are listed in Table 2.

Auto-active bait constructs were identified by stamping each transformed strain onto -His trays without previous mating. Auto active bait constructs were excluded from the analyses. A 384-format prey strain matrix (**Figure S2**) was assembled from 3-4 replicas of prey constructs stamped out on -Leu trays and grown for three – four days before mating. Bait strains were grown in liquid culture (15 – 20 hours, 30°C, 150 rpm). The prey matrix was transferred into the liquid bait culture, mixed and transferred to YPDA for mating. Mating was performed on medium for 20 – 28 hours. Colonies were then transferred directly to -His/-Trp/-Leu/-Ura/-Ade medium for protein interaction selection. Colonies were incubated over seven days and images were taken daily from day four on. Colonies were counted with a growth score of 1 or 3, 1 typically being a single small colony (background) and 3 a fully grown yeast spot. Furthermore, we excluded two prey strains from the analyses which grew repeatedly with a very high number of independently mated bait constructs (preyauto-activity). We then counted the colonies with a growth score of 3, combined technical replicates and built a ratio with the number of tests performed with the given Y2H construct combination, and got a score with a maximum of 1 for each vector combination in each biological replicate. If all biological replicates showed the interaction, we went on with the highest score found in a specific biological replicate. Lastly, we collapsed all vector combinations into protein-protein interactions and added the individual scores of eight possible vector combinations to achieve a maximal score of eight. This score of eight represents a protein-protein interaction which was detected in all eight vector combinations and all technical replicates of at least one biological replicate.

### Fly husbandry

Fly stocks were maintained at 25°C with 70% humidity in 12:12 h light/darkness cycles. For maintenance and experiments flies were fed on standard medium unless specified elsewise. All fly lines used in this study are listed in **Table S3**. Of note: *D. persimilis* Deadlock is missing from the complete set of Deadlock genes because embryo injections failed.

### *D. melanogaster* fly lines for transgene expression

PhiC31 integrase-mediated transgenesis via microinjection was performed as a paid service in the Department of Genetics at the University of Cambridge. Plasmids contained a 70 bp *attB* site for site specific recombination and the mini-white eye-marker gene for selection. Constructs were integrated on chr2 (*attP40*; FlyBase ID: FBti0114379) and originally balanced over *SM6a* (chr2). *w*^*+*^*/SM6a* (*w*^*+*^ from the recombinant plasmid and balanced with *CyO* as part of *SM6a*) males of the received strains were crossed to *if/CyO* (each on one allele of chr2, respectively) females and selected against *if* for single chromosome balancing. By discarding all *if*^*+*^, flies the only viable other flies carry maternal *CyO* and paternal *w*^*+*^. Of this generation only males were collected to ensure that the Y chromosome of the injected fly line is inherited. This eliminated the PhiC31 expression cassette originally expressed from the X chromosome in the injected flies. These males were then crossed again with *if*/*CyO* females before inbreeding and selection of homozygous *w*^*+*^/*w*^*+*^ flies. The exogenous CDS is expressed under the control of the *D. melanogaster rhi* promoter, a partial *nos* 5’-UTR and the *vasa* 3’-UTR. The construct is flanked by *gypsy* insulators to establish chromatin boarders. Coding sequences were tagged with 3xFLAG, 3xV5 and eGFP. Rhi, Del and Boot proteins were tagged N-terminally, Kipf proteins were tagged at the C-terminus.

### Co-IPs from FLAG-tagged proteins for mass spectrometry

Fly lines FS21, FS52, FS368, FS369, FS370, FS371, FS372 and FS373 (**Table S3**) were used for Co-IP MS sample preparation. All overexpressed recombinant proteins are tagged by 3xFLAG-3xV5 peptides and IPs targeted by FLAG. Flies were kept at 25°C with 70% humidity. Approximately 25 flies (3 – 5 days old) were placed in fresh bottles to reproduce. Parental flies were transferred to fresh bottles after 5 – 7 days to lay further eggs. After hatching of the F1 generation, flies (1 – 2 days old) were placed in cages on apple juice plates (25% (v/v) apple juice; 15 g/l agar; 25 g/l sucrose; 0.15% Nipagin) with yeast paste (saf-instant yeast was stirred in tap water until it becomes a sticky paste). Apple juice plates were exchanged twice a day to allow efficient laying of eggs. When flies were 3 – 5 days old, ovaries were dissected. All ovaries were placed immediately on ice in Ephuzzi Beadle Ringer solution (EBR; 10 mM HEPES pH 7.3; 130 mM NaCl; 5 mM KCl; 2 mM CaCl_2_). Isolated ovaries were collected for a maximum of 1 h before discarding the EBR solution and snap freezing in liquid nitrogen. Co-IPs of all samples were performed in biological quadruplicates of approximately 200 ovary pairs per replicate. All buffers and utensils were pre-chilled on ice before usage. The whole following workflow was performed at 4°C in a cold room. After thawing ovaries on ice, lysis was achieved by douncing (∼20 times) in 1 mL ovary protein lysis buffer 2 (OPLB2; 20 mM Tris-HCl pH 7.5, 150 mM NaCl, 2 mM MgCl_2_, 10% (v/v) Glycerol, 1 mM dithiothreitol* (DTT), 1 mM PefaBloc SC* (Sigma Aldrich, 11429868001), 0.2% (v/v) IGEPAL CA-630 (Sigma Aldrich, I3021) on ice until the solution is ‘smooth’ and seems homogenous. Reagents marked with ‘*’ were added immediately before usage. The lysates were centrifuged at 21,000 g for 5’ at 4°C, the fatty layer on top was carefully taken off and clear supernatants were transferred to fresh low retention tubes (Fisher Scientific, 11986955). Next, anti-FLAG M2 magnetic beads (Sigma Aldrich, M8823) were equilibrated in OPLB2 by placing them on a magnet, replacing the bead storage buffer (20 mM Tris-HCl pH 7.5, 150 mM NaCl) and washing them twice in OPLB2. Finally, beads were resuspended in 4:1 lysis buffer:storage volume. 16 µL of OPLB2 equilibrated beads (4 µL beads in storage volume per sample) were added to each sample. Protein capture was performed at 4°C while rotating for 3 h. Next, samples were washed four times with 1 mL of OPLB2. Beads were pelleted on a magnet, the supernatant was taken off, the beads were resuspended in 1 mL fresh OPLB2 and samples rotated for 5’ each. Afterwards samples were additionally rinsed for seven times with ovary lysis wash buffer (20 mM HEPES pH 7.4, 2 mM MgCl_2_, 150 mM NaCl) to get rid of detergents, glycerol, and other molecules, interfering with MS. For the first wash beads were resuspended fully and the tubes inverted. In the rinse buffer beads were allowed to accumulate at the magnet for 5’. During the next washes the beads were kept on the magnet, tubes were closed, and the rack was inverted. During wash number four the beads were resuspended fully again and transferred to fresh low binding tubes. After finishing all wash and rinse steps, the supernatant was removed thoroughly. To elute bound proteins, the beads were resuspended in 50 µL 2% sodium dodecyl sulfate by pipetting and rotated for 15 minutes at room temperature. The eluate was collected, and the elution step repeated. Both eluates were combined to yield 100 µL Co-IP protein samples. Both, the eluates, and the beads were snap frozen in liquid nitrogen and stored at -70°C until shipping them on dry ice.

### MS data acquisition and processing

IP samples (biological quadruplicates) were alkylated with 5mM TCEP, reduced with 10mM CAA, and digested with trypsin (∼1:10 (wt/wt)). Samples were purified utilizing the S-Trap micro columns (Protifi), following the manufacturer’s instructions: S-Trap™ micro spin column digestion protocol. Peptides were lyophilized, resuspended in 0.1% formic acid, and diluted to a concentration of 200 ng/µL, whereby 1 µL was used per MS-injection. Samples were analyzed on a TimsTOF Pro ion mobility mass spectrometer (Bruker) in-line with UltiMate 3000 UHPLC system (Thermo). Peptides were separated on a reversed-phase C18 Aurora column (25 cm × 75 µm) with an integrated Captive Spray Emitter (IonOpticks). Mobile phases A 0.1vol% formic acid in water and B 0.1vol% formic acid in ACN (Fisher Scientific, 10799704) with a flow rate of 300 nl/min, respectively. Fraction B was linearly increased from 2% to 25% in a 90 minute-gradient, increased to 40% for 10min, and a further increase to 80% for 10min, followed by re-equilibration. The spectra were recorded in diaPASEF mode^89^. Samples were recorded in technical replicates.

Subsequently, the in DIA mode recorded spectra were quantified with DIA-NN v1.8.2beta27 (ref. 90) using a synthetic library built from reviewed and selected unreviewed protein sequences from the Uni-Prot *Drosophila melanogaster* (fasta download: 04.03.2024). The protein database included transgenes and unreviewed proteins of the piRNA core complex. Expression values were log_2_-transformed, normalized across all protein groups of all samples, and averaged across their technical replicates. Further analysis only considered protein groups identified in both technical replicates. eGFP ratios for proteins of interest, were calculated by normalization to the average relative GFP abundance within each sample group. Data were visualized using R^91^, associated packages ggplot2, and VolcaNoseR^92^. The mass spectrometry data are deposited via the ProteomeXchange PRIDE partner repository with the dataset identifier PXD053049.

Briefly, in the proteomics dataset, DIA acquisition revealed quantitative data for 523 distinct protein groups, whereby 517 were detected in both technical replicates. Approximately half of the detected protein groups (292 out of 517) were detected in every sample group, and 166 protein groups are present in each sample. Conversely, 44% of the protein group abundance values (225 out of 517) displayed varying occurrences across the indicated sample groups.

### Fixed immunofluorescence of ovaries

Flies used are listed in **Table S1**. One – two days old flies were sorted (5 males 20 females) and transferred to cages on apple juice plates with yeast paste. Flies were cultivated like this at 25°C for two – three days. Apple juice plates were exchanged twice a day to allow efficient laying of eggs. Ovaries of three – five-day old female flies were then dissected into 200 µL EBR on ice. After a maximum of 30’ on ice, ovaries were fixed for 15 minutes at room temperature while rotating after addition of 1V (200 µL) of 4% paraformaldehyde (final concentration 2%). Next, ovaries were washed 3 times for 5 minutes at room temperature each in 750 µL PBX under rotation (PBX: PBS with 0.3% Triton X-100). During the second wash ovaries were dissociated into ovarioles by pipetting ∼10 times with a 200 µL tip. After washing, ovarioles were blocked in 750 µL BBX (1% (w/v) Bovine Serum Albumin in PBX) for 2 hours at room temperature while rotating. When using fluorescent antibodies, the samples were covered in Alu-foil. Ovarioles were rotating in BBX with primary antibodies at 4°C over night. On the next day, ovarioles were washed four times with 750 µL PBX at room temperature while rotating. Next, ovarioles were incubated in secondary antibody for 2 h and washed four times as before. During the second wash step DAPI (VWR, 40009) was added to a final concentration of 0.5 µg/ml. Lastly, ovarioles were mounted on slides with mounting medium (Thermo Fisher, P36961), covered with cover slips and sealed with nail polish. Mounting medium was allowed to harden for 24 hours at room temperature in the dark before microscopy. For long term storage slides were kept at 4°C in the dark. Images were acquired with a

Zeiss Axio LSM980 in Airyscan2 mode for speed (General settings: Lasers: 488 nm 5.0% intensity for Deadlocks and 3.5% intensity for Bootleggers, 850V master gain; 543 nm 2.0% intensity, 810 V master gain; 405 nm 0.5% intensity, 808 V master gain). All images were processed by the software’s Airyscan2 processing in batch mode with default settings (‘Standard’ strength). Acquired images were adjusted (Contrast, Brightness, Maximum, Minimum) in Fiji ImageJ (Schneider, Rasband, and Eliceiri 2012).

### ReLo protein-protein interaction assays

The experiments were performed as described in^58^. Briefly, we used pAc5.1-OST4-mEGFP-FspAI (pAc5.1-OST4-mEGFP; JK-268; (ref. 57)) as the basis for the bait vectors and pAc5.1-FspAI-mCherry-Daed-IsoA237-279 (pAc5.1-mCherry-MLS; JM-292) as the basis for prey vectors. OST4 is described to anchor tagged proteins to the ER^93^. The piRNA processing factor Deadalus is anchored to the outer mitochondrial membrane^94^ via a transmembrane domain (Daed^TMM^) comprising amino acid residues 237-279 (ref. 95), which was therefore used as an outer mitochondrial membrane anchor. The pAc5.1-mCherry-MLS plasmid was generated by inserting the Daed MLS (aa 237-279) into the EcoRV site of the pAc5.1-mCherry vector (T7-MJ; ref. 96) with subsequent insertion of a unique FspAI blunt-end restriction site 5’ of the mCherry coding sequence. Backbones were cut using FspAI and inserts ligated using Gibson reactions. For prey constructs 5’-agagaccccggatcgggtGCCAAC-3’ was used as overhang on the forward primer (capital letters are the added Kozak sequence) and 5’-ttggtaccgccgctggatgc-3’ was used as overhang on the reverse primer. For bait constructs 5’-gcttgaattctgcaacgtgc-3’ was used as overhang for the forward primer and 5’-gcatgagatatccagcacag-3’ was used as overhang for the reverse primer. S2R+ cells were split to a density of 10^6^ cells/mL two days before transfection. Cells were rinsed off, resuspended, and counted. 600.000 S2R+ cells were seeded on µL-Slides 4 Well IbiTreat slides (Ramcon; 80426) in 600 µL medium and co-transfected 2 hours after seeding with 300 ng per plasmid (600 ng total) by using the jetOPTIMUS transfection kit (VWR; 101000051) according to the instruction manual. Cells were imaged 30 - 48 hours post transfection without fixation with a Nikon AX-CLSM (Heidelberg, Germany) or Zeiss Axio LSM980 (Aarhus, Denmark) in confocal imaging mode.

## Notes

https://www.ebi.ac.uk/pride/archive/projects/PXD053049

### Competing Interest Statement

The authors have declared no competing interest.

### Summary of Updates

Figure 4 legend revised; minor changes in Figure 5; minor changes to the abstract.

## REFERENCES

1. Moutinho, A.F., Trancoso, F.F., and Dutheil, J.Y. (2019). The impact of protein architecture on adaptive evolution. Mol. Biol. Evol. 36, 2013–2028.

2. Afanasyeva, A., Bockwoldt, M., Cooney, C.R., Heiland, I., and Gossmann, T.I. (2018). Human long intrinsically disordered protein regions are frequent targets of positive selection. Genome Res. 28, 975–982.

3. Bricout, R., Weil, D., Stroebel, D., Genovesio, A., and Roest Crollius, H. (2023). Evolution is not Uniform Along Coding Sequences. Mol. Biol. Evol. 40. 10.1093/molbev/msad042.

4. Pare, H.K., Nguyen, A.L., Pankey, M.S., Cheeseman, I.M., and Plachetzki, D.C. (2024). Structural constraints and drivers of molecular evolution in a macromolecular complex; the kinetochore. bioRxiv, 2024.07.10.602950. 10.1101/2024.07.10.602950.

5. Peng, J., Svetec, N., and Zhao, L. (2022). Intermolecular interactions drive protein adaptive and coadaptive evolution at both species and population levels. Mol. Biol. Evol. 39. 10.1093/molbev/msab350.

6. Das, J., Vo, T.V., Wei, X., Mellor, J.C., Tong, V., Degatano, A.G., Wang, X., Wang, L., Cordero, N.A., Kruer-Zerhusen, N., et al. (2013). Cross-species protein interactome mapping reveals species-specific wiring of stress response pathways. Sci. Signal. 6, ra38.

7. Zhong, Q., Pevzner, S.J., Hao, T., Wang, Y., Mosca, R., Menche, J., Taipale, M., Taşan, M., Fan, C., Yang, X., et al. (2016). An inter-species protein-protein interaction network across vast evolutionary distance. Mol. Syst. Biol. 12, 865.

8. Lau, N.C., Seto, A.G., Kim, J., Kuramochi-Miyagawa, S., Nakano, T., Bartel, D.P., and Kingston, R.E. (2006). Characterization of the piRNA complex from rat testes. Science 313, 363–367.

9. Watanabe, T., Takeda, A., Tsukiyama, T., Mise, K., Okuno, T., Sasaki, H., Minami, N., and Imai, H. (2006). Identification and characterization of two novel classes of small RNAs in the mouse germline: retrotransposon-derived siRNAs in oocytes and germline small RNAs in testes. Genes Dev. 20, 1732–1743.

10. Aravin, A.A., Sachidanandam, R., Girard, A., Fejes-Toth, K., and Hannon, G.J. (2007). Developmentally regulated piRNA clusters implicate MILI in transposon control. Science 316, 744–747.

11. Girard, A., Sachidanandam, R., Hannon, G.J., and Carmell, M.A. (2006). A germline-specific class of small RNAs binds mammalian Piwi proteins. Nature 442, 199–202.

12. Grivna, S.T., Beyret, E., Wang, Z., and Lin, H. (2006). A novel class of small RNAs in mouse spermatogenic cells. Genes Dev. 20, 1709–1714.

13. Brennecke, J., Aravin, A.A., Stark, A., Dus, M., Kellis, M., Sachidanandam, R., and Hannon, G.J. (2007). Discrete small RNA-generating loci as master regulators of transposon activity in Drosophila. Cell 128, 1089–1103.

14. Palmer, W.H., Hadfield, J.D., and Obbard, D.J. (2018). RNA-Interference Pathways Display High Rates of Adaptive Protein Evolution in Multiple Invertebrates. Genetics 208, 1585–1599.

15. Yi, M., Chen, F., Luo, M., Cheng, Y., Zhao, H., Cheng, H., and Zhou, R. (2014). Rapid evolution of piRNA pathway in the teleost fish: implication for an adaptation to transposon diversity. Genome Biol. Evol. 6, 1393–1407.

16. Vermaak, D., Henikoff, S., and Malik, H.S. (2005). Positive selection drives the evolution of rhino, a member of the heterochromatin protein 1 family in Drosophila. PLoS Genet. 1, 96–108.

17. Kolaczkowski, B., Hupalo, D.N., and Kern, A.D. (2011). Recurrent adaptation in RNA interference genes across the Drosophila phylogeny. Mol. Biol. Evol. 28, 1033–1042.

18. Lee, Y.C.G., and Langley, C.H. (2012). Long-term and short-term evolutionary impacts of transposable elements on Drosophila. Genetics 192, 1411–1432.

19. Obbard, D.J., Gordon, K.H.J., Buck, A.H., and Jiggins, F.M. (2009). The evolution of RNAi as a defence against viruses and transposable elements. Philos. Trans. R. Soc. Lond. B Biol. Sci. 364, 99–115.

20. Simkin, A., Wong, A., Poh, Y.-P., Theurkauf, W.E., and Jensen, J.D. (2013). Recurrent and recent selective sweeps in the piRNA pathway. Evolution 67, 1081–1090.

21. Chary, S., and Hayashi, R. (2023). The absence of core piRNA biogenesis factors does not impact efficient transposon silencing in Drosophila. PLoS Biol. 21, e3002099.

22. Blumenstiel, J.P., Erwin, A.A., and Hemmer, L.W. (2016). What Drives Positive Selection in the Drosophila piRNA Machinery? The Genomic Autoimmunity Hypothesis. Yale J. Biol. Med. 89, 499–512.

23. Klattenhoff, C., Xi, H., Li, C., Lee, S., Xu, J., Khurana, J.S., Zhang, F., Schultz, N., Koppetsch, B.S., Nowosielska, A., et al. (2009). The Drosophila HP1 homolog Rhino is required for transposon silencing and piRNA production by dual-strand clusters. Cell 138, 1137–1149.

24. Mohn, F., Sienski, G., Handler, D., and Brennecke, J. (2014). The rhino-deadlock-cutoff complex licenses noncanonical transcription of dual-strand piRNA clusters in Drosophila. Cell 157, 1364–1379.

25. Le Thomas, A., Stuwe, E., Li, S., Du, J., Marinov, G., Rozhkov, N., Chen, Y.-C.A., Luo, Y., Sachidanandam, R., Toth, K.F., et al. (2014). Transgenerationally inherited piRNAs trigger piRNA biogenesis by changing the chromatin of piRNA clusters and inducing precursor processing. Genes Dev. 28, 1667–1680.

26. Akkouche, A., Kneuss, E., Bornelöv, S., Renaud, Y., Eastwood, E.L., van Lopik, J., Gueguen, N., Jiang, M., Creixell, P., Maupetit-Mehouas, S., et al. (2024). A dual histone code specifies the binding of heterochromatin protein Rhino to a subset of piRNA source loci. bioRxiv, 2024.01.11.575256. 10.1101/2024.01.11.575256.

27. Baumgartner, L., Handler, D., Platzer, S.W., Yu, C., Duchek, P., and Brennecke, J. (2022). The Drosophila ZAD zinc finger protein Kipferl guides Rhino to piRNA clusters. Elife 11. 10.7554/eLife.80067.

28. Andersen, P.R., Tirian, L., Vunjak, M., and Brennecke, J. (2017). A heterochromatin-dependent transcription machinery drives piRNA expression. Nature 549, 54–59.

29. Parhad, S.S., Yu, T., Zhang, G., Rice, N.P., Weng, Z., and Theurkauf, W.E. (2019). Adaptive evolution targets a piRNA precursor transcription network. Cell Rep. 30, 2672–2685.e5.

30. Pane, A., Jiang, P., Zhao, D.Y., Singh, M., and Schüpbach, T. (2011). The Cutoff protein regulates piRNA cluster expression and piRNA production in the Drosophila germline. EMBO J. 30, 4601–4615.

31. Zhang, Z., Wang, J., Schultz, N., Zhang, F., Parhad, S.S., Tu, S., Vreven, T., Zamore, P.D., Weng, Z., and Theurkauf, W.E. (2014). The HP1 homolog rhino anchors a nuclear complex that suppresses piRNA precursor splicing. Cell 157, 1353–1363.

32. Chen, Y.-C.A., Stuwe, E., Luo, Y., Ninova, M., Le Thomas, A., Rozhavskaya, E., Li, S., Vempati, S., Laver, J.D., Patel, D.J., et al. (2016). Cutoff Suppresses RNA Polymerase II Termination to Ensure Expression of piRNA Precursors. Mol. Cell 63, 97–109.

33. Zhang, F., Wang, J., Xu, J., Zhang, Z., Koppetsch, B.S., Schultz, N., Vreven, T., Meignin, C., Davis, I., Zamore, P.D., et al. (2012). UAP56 couples piRNA clusters to the perinuclear transposon silencing machinery. Cell 151, 871–884.

34. Hur, J.K., Luo, Y., Moon, S., Ninova, M., Marinov, G.K., Chung, Y.D., and Aravin, A.A. (2016). Splicing-independent loading of TREX on nascent RNA is required for efficient expression of dual-strand piRNA clusters in Drosophila. Genes Dev. 30, 840–855.

35. Kneuss, E., Munafò, M., Eastwood, E.L., Deumer, U.-S., Preall, J.B., Hannon, G.J., and Czech, B. (2019). Specialization of the Drosophila nuclear export family protein Nxf3 for piRNA precursor export. Genes Dev. 33, 1208–1220.

36. ElMaghraby, M.F., Andersen, P.R., Pühringer, F., Hohmann, U., Meixner, K., Lendl, T., Tirian, L., and Brennecke, J. (2019). A Heterochromatin-Specific RNA Export Pathway Facilitates piRNA Production. Cell 178, 964–979.e20.

37. Kelleher, E.S., Edelman, N.B., and Barbash, D.A. (2012). Drosophila interspecific hybrids phenocopy piRNA-pathway mutants. PLoS Biol. 10, e1001428.

38. Parhad, S.S., Tu, S., Weng, Z., and Theurkauf, W.E. (2017). Adaptive Evolution Leads to Cross-Species Incompatibility in the piRNA Transposon Silencing Machinery. Dev. Cell 43, 60–70.e5.

39. Yu, B., Lin, Y.A., Parhad, S.S., Jin, Z., Ma, J., Theurkauf, W.E., Zhang, Z.Z., and Huang, Y. (2018). Structural insights into Rhino-Deadlock complex for germline piRNA cluster specification. EMBO Rep. 19. 10.15252/embr.201745418.

40. Wang, Y.-L., Duttke, S.H.C., Chen, K., Johnston, J., Kassavetis, G.A., Zeitlinger, J., and Kadonaga, J.T. (2014). TRF2, but not TBP, mediates the transcription of ribosomal protein genes. Genes Dev. 28, 1550–1555.

41. Serebreni, L., Pleyer, L.-M., Haberle, V., Hendy, O., Vlasova, A., Loubiere, V., Nemčko, F., Bergauer, K., Roitinger, E., Mechtler, K., et al. (2023). Functionally distinct promoter classes initiate transcription via different mechanisms reflected in focused versus dispersed initiation patterns. EMBO J. n/a, e113519.

42. Russo, C.A., Takezaki, N., and Nei, M. (1995). Molecular phylogeny and divergence times of drosophilid species. Mol. Biol. Evol. 12, 391–404.

43. Shah, P.S., Wojcechowskyj, J.A., Eckhardt, M., and Krogan, N.J. (2015). Comparative mapping of host-pathogen protein-protein interactions. Curr. Opin. Microbiol. 27, 62–68.

44. Goodacre, N., Devkota, P., Bae, E., Wuchty, S., and Uetz, P. (2020). Protein-protein interactions of human viruses. Semin. Cell Dev. Biol. 99, 31–39.

45. Ghadie, M., and Xia, Y. (2019). Estimating dispensable content in the human interactome. Nat. Commun. 10, 3205.

46. Apelt, L., Knockenhauer, K.E., Leksa, N.C., Benlasfer, N., Schwartz, T.U., and Stelzl, U. (2016). Systematic protein-protein interaction analysis reveals intersubcomplex contacts in the nuclear pore complex. Mol. Cell. Proteomics 15, 2594–2606.

47. Venkatesan, K., Rual, J.-F., Vazquez, A., Stelzl, U., Lemmens, I., Hirozane-Kishikawa, T., Hao, T., Zenkner, M., Xin, X., Goh, K.-I., et al. (2009). An empirical framework for binary interactome mapping. Nat. Methods 6, 83–90.

48. Stelzl, U. (2014). E. coli network upgrade. Nat. Biotechnol. 32, 241–243.

49. Woodsmith, J., Apelt, L., Casado-Medrano, V., Özkan, Z., Timmermann, B., and Stelzl, U. (2017). Protein interaction perturbation profiling at amino-acid resolution. Nat. Methods 14, 1213–1221.

50. Bhambhani, C., Chang, J.L., Akey, D.L., and Cadigan, K.M. (2011). The oligomeric state of CtBP determines its role as a transcriptional co-activator and co-repressor of Wingless targets: CtBP oligomeric state regulates Wingless targets. EMBO J. 30, 2031–2043.

51. Zhao, R., Shen, J., Green, M.R., MacMorris, M., and Blumenthal, T. (2004). Crystal structure of UAP56, a DExD/H-box protein involved in pre-mRNA splicing and mRNA export. Structure 12, 1373–1381.

52. Obbard, D.J., Maclennan, J., Kim, K.-W., Rambaut, A., O’Grady, P.M., and Jiggins, F.M. (2012). Estimating divergence dates and substitution rates in the Drosophila phylogeny. Mol. Biol. Evol. 29, 3459–3473.

53. Müller, J., Hart, C.M., Francis, N.J., Vargas, M.L., Sengupta, A., Wild, B., Miller, E.L., O’Connor, M.B., Kingston, R.E., and Simon, J.A. (2002). Histone methyltransferase activity of a Drosophila Polycomb group repressor complex. Cell 111, 197–208.

54. Cao, R., Wang, L., Wang, H., Xia, L., Erdjument-Bromage, H., Tempst, P., Jones, R.S., and Zhang, Y. (2002). Role of histone H3 lysine 27 methylation in Polycomb-group silencing. Science 298, 1039–1043.

55. Czermin, B., Melfi, R., McCabe, D., Seitz, V., Imhof, A., and Pirrotta, V. (2002). Drosophila enhancer of Zeste/ESC complexes have a histone H3 methyltransferase activity that marks chromosomal Polycomb sites. Cell 111, 185–196.

56. Pekovic, F., Rammelt, C., Kubíková, J., Metz, J., Jeske, M., and Wahle, E. (2023). RNA binding proteins Smaug and Cup induce CCR4-NOT-dependent deadenylation of the nanos mRNA in a reconstituted system. Nucleic Acids Res. 51, 3950–3970.

57. Kubíková, J., Ubartaitė, G., Metz, J., and Jeske, M. (2023). Structural basis for binding of Drosophila Smaug to the GPCR Smoothened and to the germline inducer Oskar. Proc. Natl. Acad. Sci. U. S. A. 120, e2304385120.

58. Salgania, H.K., Metz, J., and Jeske, M. (2024). ReLo is a simple and rapid colocalization assay to identify and characterize direct protein-protein interactions. Nat. Commun. 15, 2875.

59. Sewalt, R.G., Gunster, M.J., van der Vlag, J., Satijn, D.P., and Otte, A.P. (1999). C-Terminal binding protein is a transcriptional repressor that interacts with a specific class of vertebrate Polycomb proteins. Mol. Cell. Biol. 19, 777–787.

60. Turner, J., and Crossley, M. (2001). The CtBP family: enigmatic and enzymatic transcriptional co-repressors. Bioessays 23, 683–690.

61. Nielsen, R., Bustamante, C., Clark, A.G., Glanowski, S., Sackton, T.B., Hubisz, M.J., Fledel-Alon, A., Tanenbaum, D.M., Civello, D., White, T.J., et al. (2005). A scan for positively selected genes in the genomes of humans and chimpanzees. PLoS Biol. 3, e170.

62. Sackton, T.B., Lazzaro, B.P., Schlenke, T.A., Evans, J.D., Hultmark, D., and Clark, A.G. (2007). Dynamic evolution of the innate immune system in Drosophila. Nat. Genet. 39, 1461–1468.

63. Obbard, D.J., Welch, J.J., Kim, K.-W., and Jiggins, F.M. (2009). Quantifying adaptive evolution in the Drosophila immune system. PLoS Genet. 5, e1000698.

64. Slodkowicz, G., and Goldman, N. (2020). Integrated structural and evolutionary analysis reveals common mechanisms underlying adaptive evolution in mammals. Proc. Natl. Acad. Sci. U. S. A. 117, 5977–5986.

65. Enard, D., Cai, L., Gwennap, C., and Petrov, D.A. (2016). Viruses are a dominant driver of protein adaptation in mammals. Elife 5. 10.7554/eLife.12469.

66. Zhang, J., and Yang, J.-R. (2015). Determinants of the rate of protein sequence evolution. Nat. Rev. Genet. 16, 409–420.

67. Booker, T.R., Jackson, B.C., and Keightley, P.D. (2017). Detecting positive selection in the genome. BMC Biol. 15, 98.

68. Sawyer, S.L., Wu, L.I., Emerman, M., and Malik, H.S. (2005). Positive selection of primate TRIM5alpha identifies a critical species-specific retroviral restriction domain. Proc. Natl. Acad. Sci. U. S. A. 102, 2832–2837.

69. Barrett, R.D.H., Laurent, S., Mallarino, R., Pfeifer, S.P., Xu, C.C.Y., Foll, M., Wakamatsu, K., Duke-Cohan, J.S., Jensen, J.D., and Hoekstra, H.E. (2019). Linking a mutation to survival in wild mice. Science 363, 499–504.

70. Karageorgi, M., Groen, S.C., Sumbul, F., Pelaez, J.N., Verster, K.I., Aguilar, J.M., Hastings, A.P., Bernstein, S.L., Matsunaga, T., Astourian, M., et al. (2019). Genome editing retraces the evolution of toxin resistance in the monarch butterfly. Nature 574, 409–412.

71. Estojak, J., Brent, R., and Golemis, E.A. (1995). Correlation of two-hybrid affinity data with in vitro measurements. Mol. Cell. Biol. 15, 5820–5829.

72. Nekrasov, M., Wild, B., and Müller, J. (2005). Nucleosome binding and histone methyltransferase activity of Drosophila PRC2. EMBO Rep. 6, 348–353.

73. Tie, F., Stratton, C.A., Kurzhals, R.L., and Harte, P.J. (2007). The N terminus of Drosophila ESC binds directly to histone H3 and is required for E(Z)-dependent trimethylation of H3 lysine 27. Mol. Cell. Biol. 27, 2014–2026.

74. Orr, H.A. (1998). The population genetics of adaptation: The distribution of factors fixed during adaptive evolution. Evolution 52, 935–949.

75. Moutinho, A.F., Eyre-Walker, A., and Dutheil, J.Y. (2022). Strong evidence for the adaptive walk model of gene evolution in Drosophila and Arabidopsis. PLoS Biol. 20, e3001775.

76. Wang, L., Barbash, D.A., and Kelleher, E.S. (2020). Adaptive evolution among cytoplasmic piRNA proteins leads to decreased genomic auto-immunity. PLoS Genet. 16, e1008861.

77. Kelleher, E.S., Barbash, D.A., and Blumenstiel, J.P. (2020). Taming the Turmoil Within: New Insights on the Containment of Transposable Elements. Trends Genet. 36, 474–489.

78. Kelleher, E.S. (2021). Protein-protein interactions shape genomic autoimmunity in the adaptively evolving Rhino-Deadlock-Cutoff complex. Genome Biol. Evol. 13, evab132.

79. Blumenstiel, J.P. (2025). From the cauldron of conflict: Endogenous gene regulation by piRNA and other modes of adaptation enabled by selfish transposable elements. Seminars in Cell & Developmental Biology 164, 1–12.

80. Van Valen, L. (1973). A New Evolutionary Law. Evol. Theory 1, 1–30.

81. Luo, S., Zhang, H., Duan, Y., Yao, X., Clark, A.G., and Lu, J. (2020). The evolutionary arms race between transposable elements and piRNAs in Drosophila melanogaster. BMC Evol. Biol. 20, 14.

82. Ellison, C.E., Kagda, M.S., and Cao, W. (2020). Telomeric TART elements target the piRNA machinery in Drosophila. PLoS Biol. 18, e3000689.

83. Wierzbicki, F., Kofler, R., and Signor, S. (2023). Evolutionary dynamics of piRNA clusters in Drosophila. Mol. Ecol. 32, 1306–1322.

84. Rozas, J., Ferrer-Mata, A., Sánchez-DelBarrio, J.C., Guirao-Rico, S., Librado, P., Ramos-Onsins, S.E., and Sánchez-Gracia, A. (2017). DnaSP 6: DNA Sequence Polymorphism analysis of large data sets. Mol. Biol. Evol. 34, 3299–3302.

85. Yang, Z. (2007). PAML 4: phylogenetic analysis by maximum likelihood. Mol. Biol. Evol. 24, 1586–1591.

86. Altschul, S.F., Gish, W., Miller, W., Myers, E.W., and Lipman, D.J. (1990). Basic local alignment search tool. J. Mol. Biol. 215, 403–410.

87. Hegele, A., Kamburov, A., Grossmann, A., Sourlis, C., Wowro, S., Weimann, M., Will, C.L., Pena, V., Lührmann, R., and Stelzl, U. (2012). Dynamic protein-protein interaction wiring of the human spliceosome. Mol. Cell 45, 567–580.

88. Worseck, J.M., Grossmann, A., Weimann, M., Hegele, A., and Stelzl, U. (2012). A stringent yeast two-hybrid matrix screening approach for protein-protein interaction discovery. Methods Mol. Biol. 812, 63–87.

89. Meier, F., Brunner, A.-D., Frank, M., Ha, A., Bludau, I., Voytik, E., Kaspar-Schoenefeld, S., Lubeck, M., Raether, O., Bache, N., et al. (2020). diaPASEF: parallel accumulation-serial fragmentation combined with data-independent acquisition. Nat. Methods 17, 1229–1236.

90. Demichev, V., Messner, C.B., Vernardis, S.I., Lilley, K.S., and Ralser, M. (2020). DIA-NN: neural networks and interference correction enable deep proteome coverage in high throughput. Nat. Methods 17, 41–44.

91. R Core Team (2023). R: A Language and Environment for Statistical Computing. https://www.R-project.org/.

92. Goedhart, J., and Luijsterburg, M.S. (2020). VolcaNoseR is a web app for creating, exploring, labeling and sharing volcano plots. Sci. Rep. 10, 20560.

93. Möckli, N., Deplazes, A., Hassa, P.O., Zhang, Z., Peter, M., Hottiger, M.O., Stagljar, I., and Auerbach, D. (2007). Yeast split-ubiquitin-based cytosolic screening system to detect interactions between transcriptionally active proteins. Biotechniques 42, 725–730.

94. Munafò, M., Manelli, V., Falconio, F.A., Sawle, A., Kneuss, E., Eastwood, E.L., Seah, J.W.E., Czech, B., and Hannon, G.J. (2019). Daedalus and Gasz recruit Armitage to mitochondria, bringing piRNA precursors to the biogenesis machinery. Genes Dev. 33, 844–856.

95. Salgania, H.K., Metz, J., Lingren, E., Bleischwitz, C., Hauser, D., Oliveras Máté, K., Bollack, D., Lahr, F., Garbelyanski, A., and Jeske, M. (2024). Molecular insight into the network of Drosophila cytoplasmic piRNA pathway proteins through a combination of systematic interaction screening and structural prediction. bioRxiv, 2024.05.31.596839. 10.1101/2024.05.31.596839.

96. Jeske, M., Müller, C.W., and Ephrussi, A. (2017). The LOTUS domain is a conserved DEAD-box RNA helicase regulator essential for the recruitment of Vasa to the germ plasm and nuage. Genes Dev. 31, 939–952.

97. Edgar, R.C. (2004). MUSCLE: multiple sequence alignment with high accuracy and high throughput. Nucleic Acids Res. 32, 1792–1797.

