## Supplemental figures 1-8 for "Recurrent innovation of protein-protein interactions in the *Drosophila* piRNA pathway"

### Figure S1

**A**

| Gene | Species | range of % sequence analyzed | 2ΔlogL (log ratio) M7,M8 | p value | positively selected sites (prob. > 95%)) |
|---|---|---|---|---|---|
| *Boot* | *mel, sim sec, mau, ere, yak, tei, eug, bia, suz, tak, ele, rho, fic* | 76 - 92 | 0.06 | 0.97 | NA |
| *CtBP* | *mel, sim sec, mau, ere, yak, tei, eug, bia, suz, tak, ele, rho, fic* | 77 - 97 | 0.00 | 1.00 | NA |
| *cuff* | *mel, sim sec, mau, ere, yak, tei, eug, bia, suz, tak, ele, rho, fic* | 92 - 96 | 1.28 | 0.53 | NA |
| *del* | *mel, sim sec, mau, ere, yak, tei, eug, bia, suz, tak, ele, rho, fic* | 55 - 71 | 9.85 | 0.01 | NA |
| *kipf* | *mel, sim sec, mau, ere, yak, tei, eug, suz, tak, ele, fic* | 50 -72 | 17.60 | 0.00 | 97P |
| *moon* | *mel, sim sec, mau, ere, yak, tei, eug, suz, tak, ele, rho, fic* | 69 - 98 | 13.97 | 0.00 | 164E |
| *Nxf3* | *mel, sim sec, mau, ere, yak, tei, eug, bia, suz, tak, ele, rho, fic* | 90 -94 | 1.67 | 0.43 | NA |
| *rhi* | *mel, sim sec, mau, ere, yak, tei, eug, bia, suz, tak, ele, rho, fic* | 14 - 93 | 12.26 | 0.00 | 57K, 70H, 74I |
| *TFIIA-S* | *mel, sim sec, mau, ere, yak, tei, eug, bia, suz, tak, ele, rho, fic* | 99 - 100 | 0.00 | 1.00 | NA |
| *TRF2* | *mel, sim sec, mau, ere, yak, tei, tak, rho* | 50 - 70 | 95.03 | 0.00 | *** |
| *UAP56* | *mel, sim sec, mau, ere, yak, tei, eug, bia, suz, tak, ele, rho, fic* | 100 | 0.01 | 1.00 | 1.00 |

**Figure S1. Positive selection test results of codeml analyses from PAML related to Figure 1**
Results of codeML analysis (PAML software package) of the genes involved in germline piRNA precursor biogenesis (Figure 1A). TRF2 residues under positive selection are marked with ***: 2G*, 4A**, 29F*, 54R*, 101T**, 102R*, 126N*, 157G**, 159S*, 180R*, 184S*, 200S*, 236F**, 291S*, 364F*, 367E*. Of note, all of these residues are part of the long unstructured C-terminal region of the long isoform of TRF2 and not in the short isoform.

### Figure S2

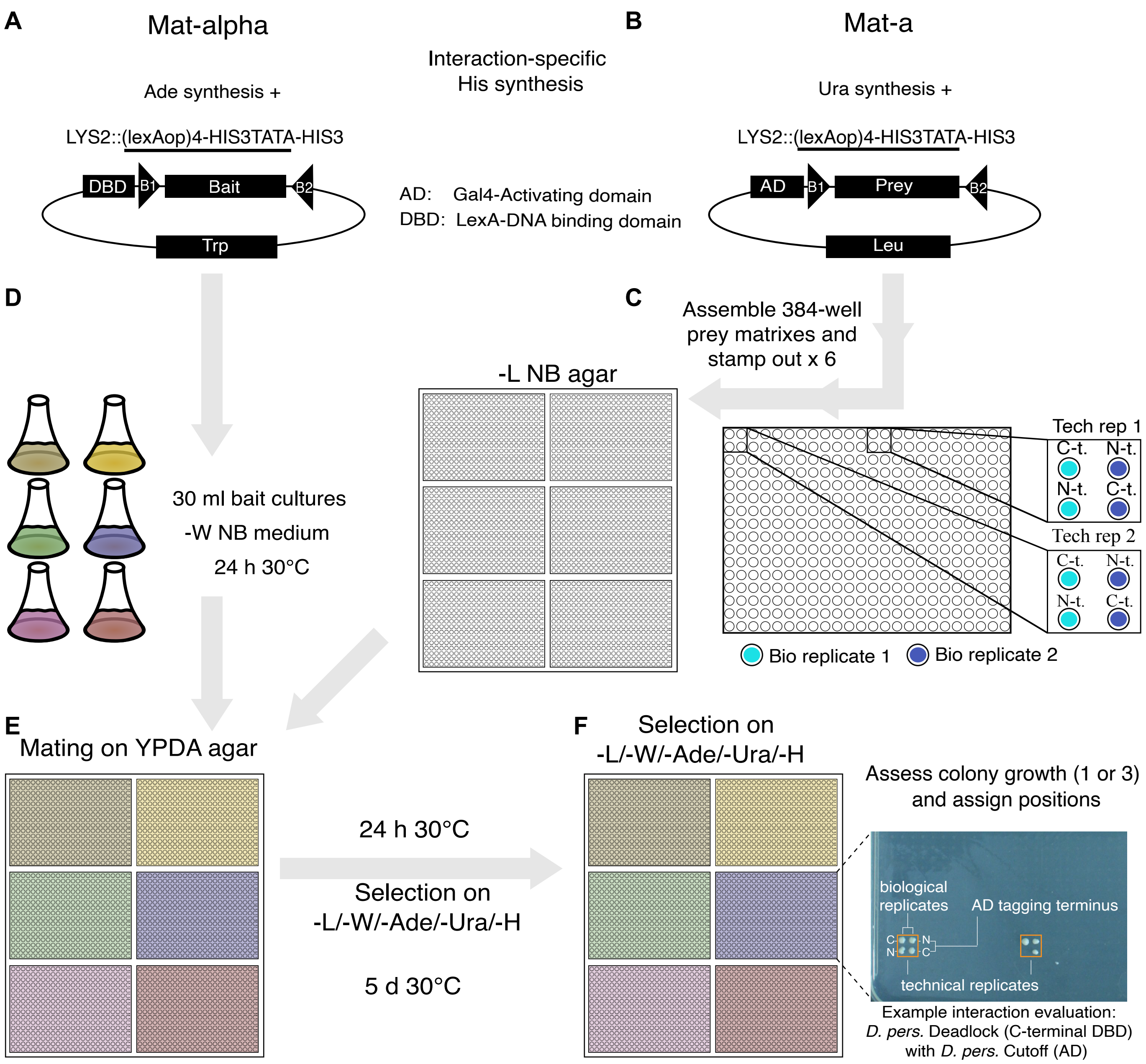


**Figure S2. Yeast-two-hybrid screen workflow**
(A-B) Mat-alpha and Mat-a strains of S. cerevisiae were transformed with bait (A) and prey (B) vectors, respectively and grown in selective medium. (C) Prey yeast strains were assembled into a fixed 384-well format stamped out in six replicates per tray and grown on selective medium. For mating, the prey matrices were stamped into liquid cultures of bait strains (D) and the mixture was transferred to non-selective YPDA medium (E). (F) After growth, the mated colonies (one bait vector per 384-format prey matrix) were stamped onto limited medium (-Leu: prey plasmid; -Trp: bait plasmid; -Ade: selection against Mat-alpha; -Ura: selection against Mat-a; -His: selection for PPI). The trays were then incubated at 30°C for four – seven days before images were taken.

### Figure S3

**A**

| | | | |
|---|---|---|---|
| ① | Clone 11 genes from 5 species, transform and mate all-against-all | 89,088 | yeast matings |
| ② | Drop autoreactive bait (10) and prey (2) and detect colony-forming matings | 4,581 | colonies |
| ③ | Aggregate colonies in technical and biological replicates | 1,573 | interactions detected |
| ④ | Drop weakly replicated interactions with weak growth | 627 | high-conf. interactions |
| ⑤ | Collapse all vectors supporting each tested protein-protein interaction (PPIs) | 263 | PPIs |

**B** **Interaction replication score calculation:**

1. Per vector pair score: colonies [Count] / Interactions tested (max 1)
2. If reproduced in biological replicates: Use the highest individual score
3. Add the highest scores of each vector combination (max 8)

| Final score: | < 0.5 | ≥ 0.5 | ≥ 0.75 | ≥ 1 | ≥ 2 | ≥ 4 |
|---|---|---|---|---|---|---|
| # of interactions: | 263 | 238 | 199 | 172 | 92 | 46 |
| → intra-species: | 64 | 55 | 46 | 42 | 29 | 14 |
| → inter-species: | 199 | 183 | 153 | 130 | 63 | 32 |

**Figure S3. Yeast-two-hybrid data evaluation and replication score calculation**
(A) Table showing the summary statistics following each step in the yeast-to-hybrid screen data evaluation workflow, resulting in the detection of 263 individual protein-protein interactions. (B) Overview of the calculation of the yeast-two-hybrid growth replication score, including summary statistics for six different score thresholds. Based on congruence with previous data describing protein-protein interactions within the tested network a threshold of ≥ 0.75 was selected for calling positive interactions in our screen.

### Figure S4

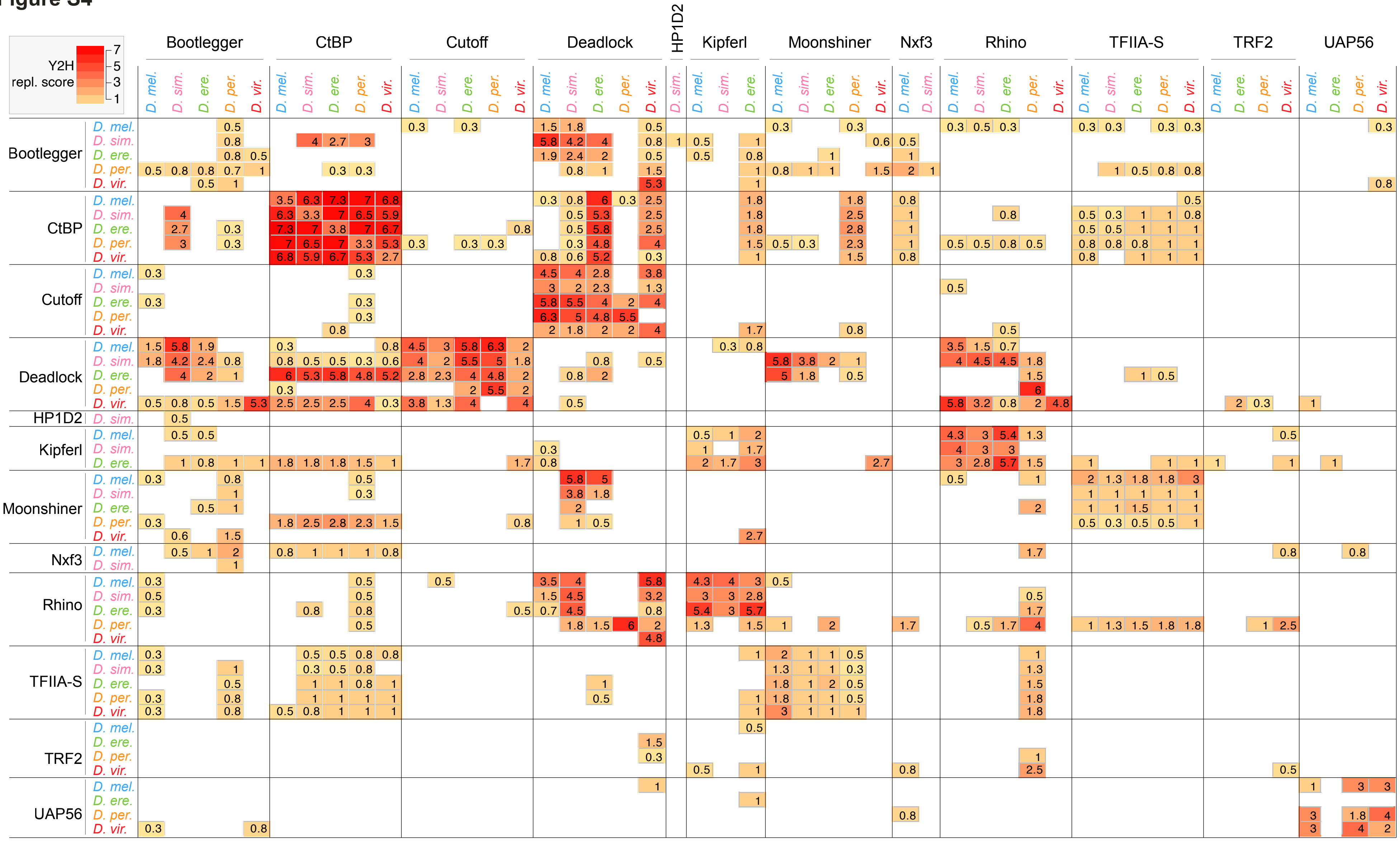


**Figure S4. Yeast-two-hybrid replication scores for all tested interactions**
Y2H replication scores are noted in the matrix as numbers as well as by heat map color code (see legend). The replication scores were calculated as described in Figure S3B.

### Figure S5

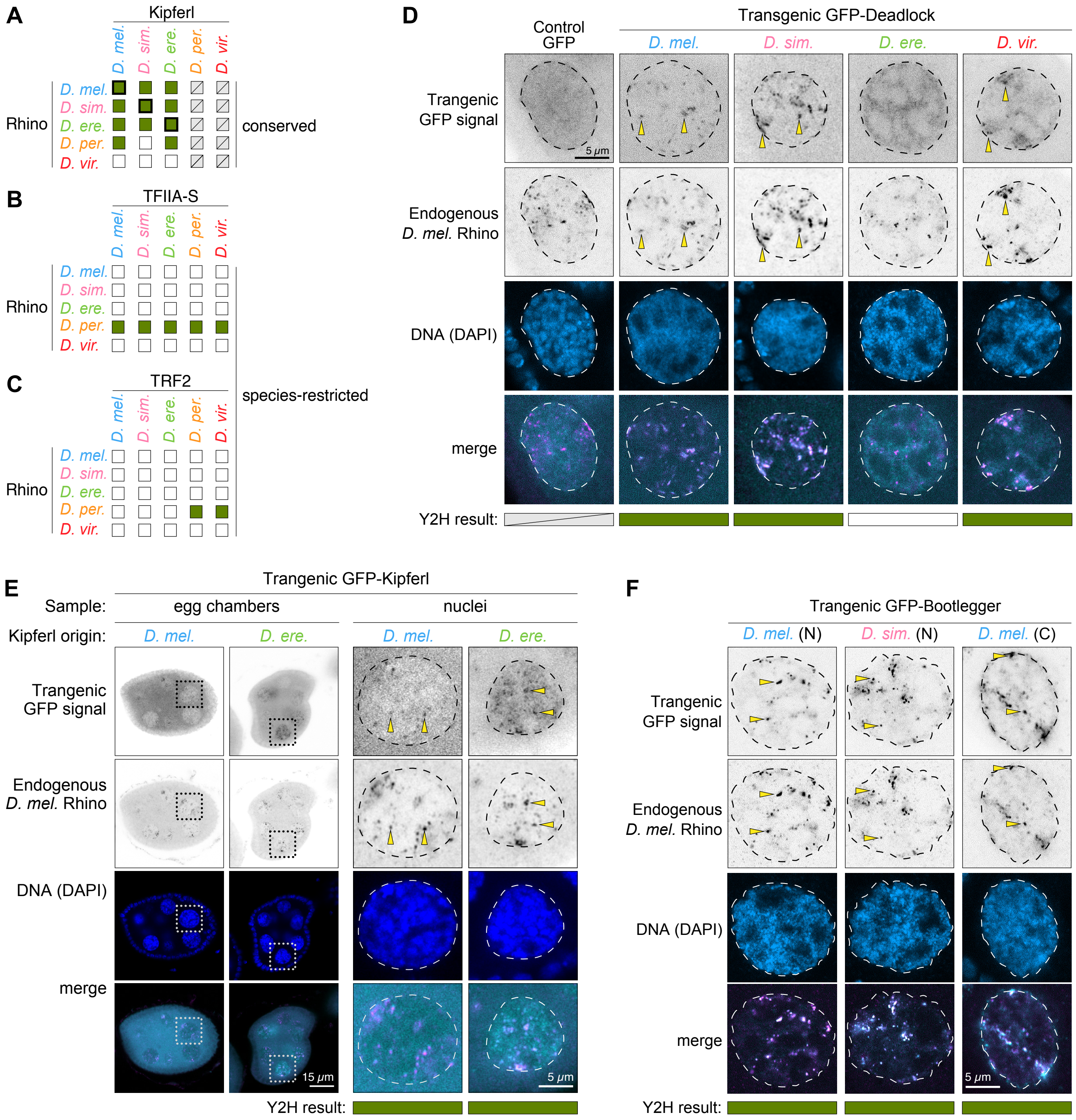


**Figure S5. Supplementary data related to Figure 3**

(A-C) Summary of interactions detected by yeast-two-hybrid between orthologs of Rhino and Kipferl (A), TFIIA-S (B), and Trf2 (C). Intra-species (thick box outline) and inter-species (thin box outline) interactions are shown as filled green boxes, while empty boxes indicate absence of interaction detection above the replication score threshold. Grey crossed-out circles for Kipferl denote absence of the gene in those species. Pattern: evolutionary signature of the interaction (see main text for details). (D-F) Confocal microscopy images showing the localization of endogenous D. melanogaster Rhino (anti-Rhi IF) and GFP-tagged transgenic Deadlock (D), Kipferl (E), and Bootlegger (F) from the indicated species. Dashed line: nuclear border as determined by DAPI staining. Yellow arrows highlight co-localizing foci of endogenous Rhino IF signal and transgenic GFP-tagged proteins.

### Figure S6

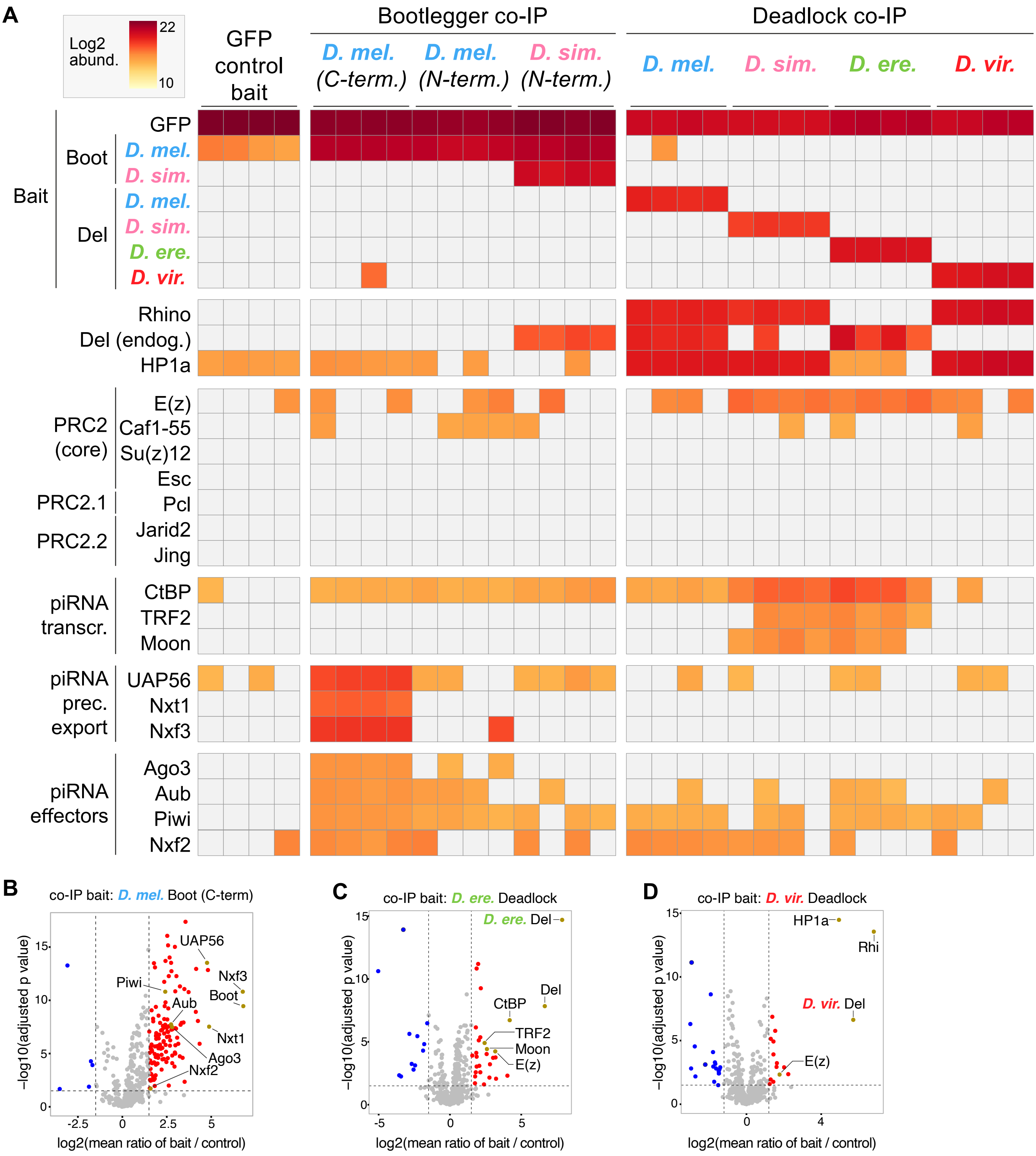


**Figure S6. Supplementary co-IP/MS data related to Figure 4**
(A) Heatmap diagram showing log2-transformed mass spectrometry abundance signal values for the proteins indicated to the left in the co-IP/MS bait samples denoted above the diagram. (B-D) Volcano plots displaying co-IP/mass spectrometry data from the co-IP baits indicated above each plot. The x-axes show log2 fold change between bait and control IP averaged over four biological replicates with each two technical replicates. The y-axes display the negative log10 of adjusted p values from t-tests for enrichment in bait compared to control co-IP. Red dots represent proteins enriched more than 1.5 fold with -log10(adjusted p values) higher than 2.

### Figure S7

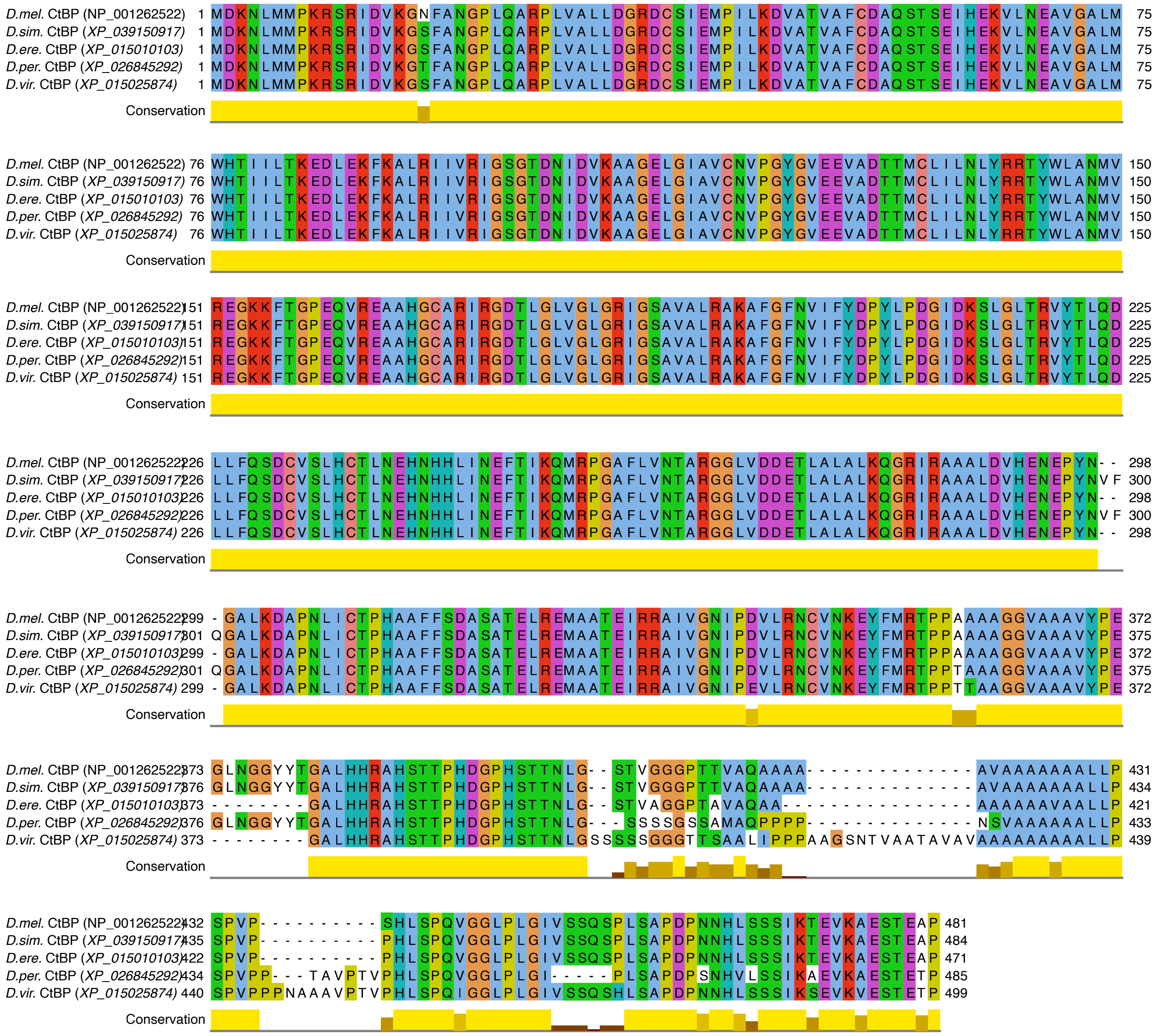


**Figure S7. Protein sequence alignment of CtBP from five Drosophila species**
The protein sequences were retrieved from NCBI using the noted accession numbers, aligned using the MUSCLE program (Edgar 2004) and displayed using Jalview. Amino acids are colored according to the 'Clustal' color scheme. Conservation is shown as bar diagrams with maximum values of 10, representing full conservation in all included species.

### Figure S8

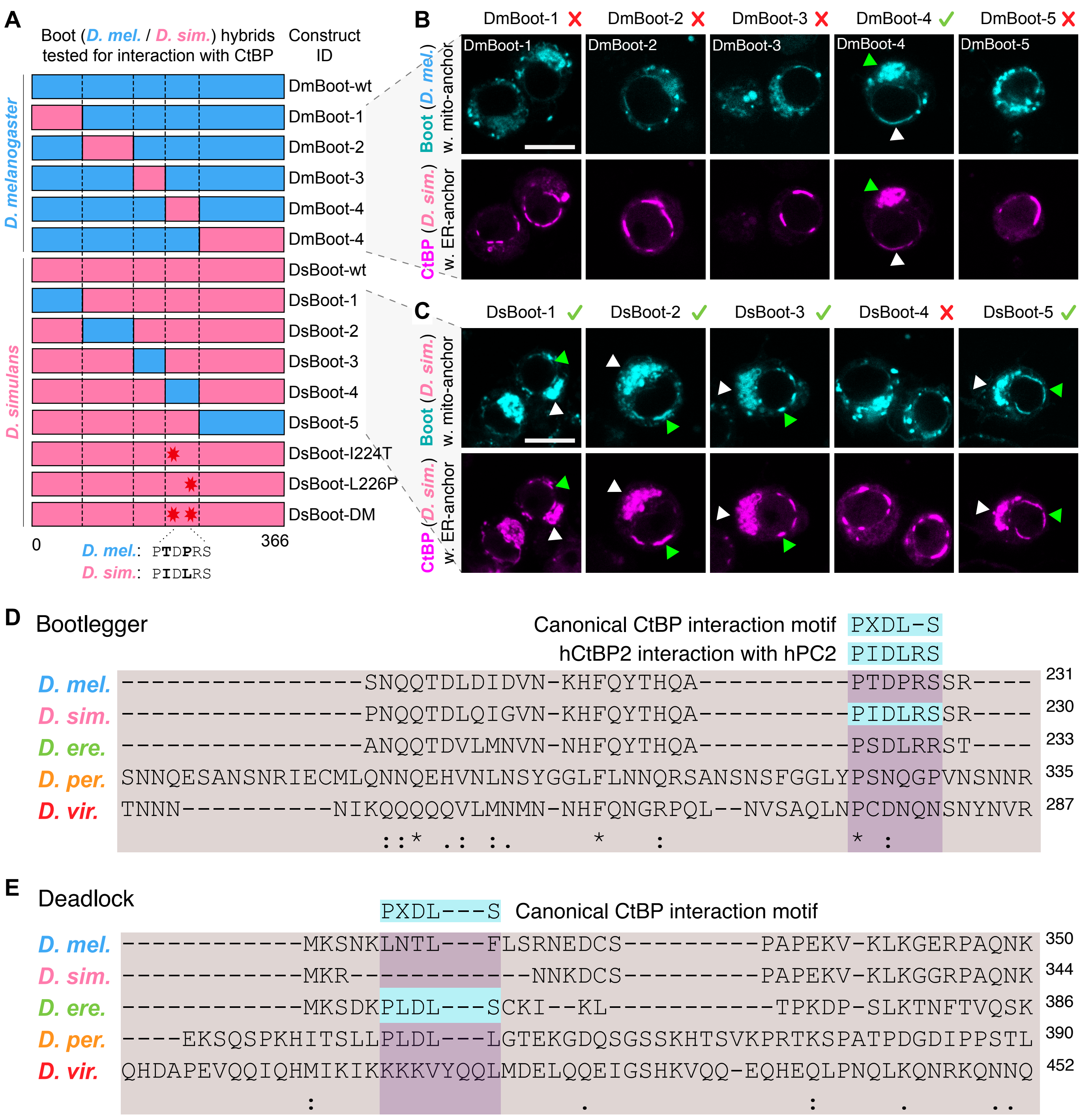


**Figure S8. Supplementary ReLo protein interaction analyses related to Figure 5**
(A) Schematic representation of domain-swap constructs between the D. melanogaster (blue) and D. simluans (pink) Bootlegger orthologs with the interaction results indicated to right right by red crosses or green check marks. Black stars indicate the tested interaction site point mutants. (B-C) Fluorescence microscopy images of ReLo protein interaction assays testing interaction between CtBP and D. melanogaster Bootlegger proteins harboring swapped domains from D. simluans (B) or vice versa (C); see also (A). Shown are representative cells from >10 images. Green arrows indicate protein accumulation in interrupted ring structures around the nucleus. White arrows show protein accumulation at cytoplasmic sites. Scale bars indicate 8 $\mu m$ in size. Red cross and green check mark denote whether protein interaction was concluded or not, respectively, based on the ReLo assays. (D-E) Amino acid sequence alignment around the identified CtBP interaction motifs in Bootlegger (D) and Deadlock (E) from the five investigated Drosophila species.
